# Sexual antagonism, mating systems, and recombination suppression on sex chromosomes

**DOI:** 10.1101/2025.05.15.654257

**Authors:** Ewan Flintham, Charles Mullon

## Abstract

The suppression of recombination between sex chromosomes is a widespread feature of genetic sex determination systems. Competing explanations for this evolution fall into two broad categories: recombination arrest driven by the capture of sexually antagonistic variation versus arrest driven without sex-specific selection. Using population-genetic models, we compare the substitution rates of full recombination suppressors (such as inversions) driven by sexually antagonistic selection to those expected under neutrality. We consider both XY and ZW systems, mating ecologies ranging from random mating to polygyny with high male reproductive variance, suppressors arising on heterogametic (Y or W) versus homogametic (X or Z) chromosomes, and the effects of deleterious alleles segregating at other loci on the chromosome. We find that even very weak sexual antagonism is sufficient to expedite suppressor substitution by orders of magnitude relative to neutrality. This acceleration is robust to the presence of deleterious variation, although high variance in male reproductive success can diminish substitution rates of Y-linked suppressors in XY systems. By contrast, in ZW systems, elevated male reproductive variance tends to favour faster recombination arrest via W-linked suppressors, leading to higher overall rates of suppression than in comparable XY systems. Despite stronger selection on Y/W-linked suppressors, recombination arrest driven by sexual antagonism is often expected to arise from homogametic (X/Z-linked) suppressors, because their mutational input is higher and they can experience weaker drift. Together, these results yield testable predictions for the genomic distribution of sex-chromosome inversions across taxa differing in mating system and sex-determination system.

## 1 Introduction

In many species with separate sexes, an individual’s sex is determined by its karyotype, in particular the identity of its sex chromosomes (Bachtrog et al., 2014). These chromosomes contain the genes that instigate sex-specific development, which are held together in a genomic region that does not recombine in at least one sex (a sex-determining region, or “SDR”, Otto et al., 2011; Wright et al., 2016). For example in XY species, sex is assigned according to the presence or absence of a dominant male-determining locus on the Y-chromosome, around which recombination is suppressed in males. This situation is mirrored in ZW species, where a dominant female-determining factor sits on the W-chromosome, leading to recombination suppression in females. Strong genetic linkage between sex-determining loci in the heterogametic sex is thought to be adaptive as it ensures alleles responsible for sexual development are co-inherited (Charlesworth and Charlesworth, 2005; Nei, 1969). However, regions of suppressed recombination that extend well beyond the vicinity of sex-determining loci, often encompassing the majority or even entirety of the chromosome, have evolved independently in many species (Bachtrog et al., 2014; Bergero and Charlesworth, 2009). Indeed, extensive recombination suppression in the heterogametic sex is considered a hallmark of a sex chromosome system (Otto, 2014).

The classical explanation for the evolution of recombination suppression on sex chromosomes is that selection favours tighter linkage between a sex-determining factor and alleles that are beneficial when expressed in that sex (i.e., that contribute to sex-specific adaptation, Beukeboom and Perrin, 2014; Fisher, 1931; Ponnikas et al., 2018; Wright et al., 2016). Theoretical population genetic models have been integral to our understanding of this process. Early models showed that selection favours the invasion of a Y-linked recombination suppressor if it captures a male SDR and the male-beneficial allele at a sexually antagonistic locus under balancing selection (modelled as a single di-allelic site with fixed fitness effects; Charlesworth and Charlesworth, 1980; Nei, 1969). Symmetrically, Charlesworth and Charlesworth (1980) also demonstrated that selection favours an X-linked suppressor if it captures the SDR-homologue and the female-beneficial allele at such a locus. Lenormand (2003) later found that a modifier decreasing recombination by a small amount would be favoured if it increased linkage between the SDR and a previously weakly linked polymorphic locus experiencing sex-specific selection – although without balancing selection, variation at such a locus would be transient, limiting the scope for selection on recombination. This result was then found to hold for most levels of linkage between a modifier, the SDR and a locus under selection (Otto, 2014). Finally, Olito and Abbott (2025) and Olito et al. (2024) estimated the invasion probability of an inversion linking a Y-linked SDR with a di-allelic sexually antagonistic locus with weak fitness effects in a large well-mixed population, and used this to make predictions about the chromosomal length affected by recombination suppression.

In contrast to the classic sexual antagonism explanation, a more recent group of models have shown that it is possible for recombination suppression on sex chromosomes to evolve without sex-specific selection (reviewed in Charlesworth and Olito, 2024; Jay et al., 2024; Olito and Abbott, 2025; Ponnikas et al., 2018). Here, recombination suppressors capture the SDR and a nearby region of the chromosome that is either (i) neutral (i.e., contains no segregating loci affecting fitness; Charlesworth and Charlesworth, 2005; Ironside, 2010; Jeffries et al., 2021) and then fixes by chance owing to genetic drift; or (ii) includes loci at which deleterious recessive variants segregate in the population. In this second case, a “lucky” suppressor mutant that happens to appear in a particularly unloaded background (i.e., captures a set of loci at which fewer than average deleterious alleles are present; Lenormand and Roze, 2022), experiences a transient fitness advantage that helps drive its fixation (Charlesworth and Olito, 2024; Jay et al., 2024; Olito and Abbott, 2025; Olito et al., 2024), although the fixation probabilities of these suppressors generally remain well below those of neutral alleles (Roze and Lenormand, 2025). In both scenarios, recombination arrest must later be stabilised in the face of mutations that restore recombination (e.g., reversions; Lenormand and Roze, 2024). This may occur through the development of structural incompatibilities between sex chromosome homologues after recombination arrest (Jay et al., 2024), or through sex-specific divergence in gene regulation at loci within the non-recombining region, leading to low fitness recombinants (Lenormand and Roze, 2022).

While theory has demonstrated that sexual antagonism and the fixation of neutral or lucky suppressors can in principle drive recombination arrest, the relative contribution of these mechanisms to sex chromosome evolution, as well as the genetic signatures they may be associated with, remain understudied (Jay et al., 2024; Lenormand and Roze, 2024; Olito and Abbott, 2025; Ponnikas et al., 2018). In particular, it is not known how strongly sexual antagonism, genetic drift, and selection on deleterious variation affect the substitution of recombination suppressors, and whether they do so similarly for modifiers arising in different sex-linked regions. For example, sexually antagonistic polymorphisms generate stronger positive selection on full suppressors arising on the heterogametic (Y or W) than on homogametic (X or Z) sex chromosomes (Blackmon and Brandvain, 2017; Charlesworth and Charlesworth, 1980), but the impact of this asymmetry on evolutionary rates – once mutation and drift are taken into account – remains underexplored. Meanwhile, studies considering lucky inversions have predominantly focused on fixation probabilities in autosomes and Y-chromosomes (Charlesworth and Olito, 2024; Jay et al., 2024; Roze and Lenormand, 2025), whereas substitution rates on homogametic chromosomes are yet to be quantified. Consequently, we lack predictions for whether recombination evolution under different models should occur on comparable timescales, or involve modifiers with similar genomic distributions. Yet with the increasing availability of large cross-taxa genomic datasets (e.g., Jeffries et al., 2025), such predictions may prove useful in delineating the mechanisms driving sex chromosome evolution by providing tests for comparative studies.

Characterising the tempo of recombination evolution on sex chromosomes, however, is not straight-forward. Substitution rate of recombination modifiers depend on multiple factors, including the nature of segregating sexually antagonistic and deleterious variation, as well as the strength of genetic drift experienced by sex-linked genomic regions and so on demographic parameters such as the mating system (Laporte and Charlesworth, 2002; Vicoso and Charlesworth, 2009). Indeed, by modulating the effective population sizes of sex-linked genomic regions, mating system variation may have especially strong effects on the relative rates of recombination evolution along different sex chromosomes (e.g., X vs Y), as well as the pace of recombination suppression in different sex-determination systems (e.g., in XY vs ZW systems). However, the consequences of these effects for suppressor substitution rates across X, Y, Z, and W chromosomes have yet to be characterised or integrated into existing theory on sex chromosome evolution.

Here, we address some of these issues by analysing a series of mathematical and computational models of recombination suppression to better understand the tempo of sex chromosome evolution. We do this under different assumptions about the segregation of sexually antagonistic and universally deleterious alleles, as well as the mating and sex-determination systems. Specifically, we first characterise the nature of sexually antagonistic variation likely to evolve on the PAR. Starting with the Y-chromosome as a basis, we then investigate the strength of selection for recombination suppression driven by sexual antagonism by calculating the fixation probability and substitution rate of a full recombination suppressor capturing a sex-specifically selected locus. We then extend our analyses to consider fixation probabilities for recombination suppressors appearing in different sex-linked regions (X, Y, Z, and W chromosomes) and under different mating systems. Finally, we use individual-based simulations to estimate fixation probabilities for recombination suppressors that can capture a sexually antagonistic locus at sites at which ubiquitously deleterious alleles segregate.

## 2 Model

### 2.1 Lifecycle and fitness

We consider a diploid population of large and constant size *N*, composed of equal numbers of adult males and females, with non-overlapping generations. Males and females reproduce to generate zygotes whose sex is genetically determined such that the primary sex ratio is unbiased. Adults die and juveniles compete sex-specifically to become the male and female adults of the next generation. As a baseline, reproduction occurs through the random union of male and female gametes, but later we incorporate a more flexible mating model that allows for male-biased reproductive variance.

Each individual expresses a quantitative trait *z* that influences its reproductive success in a sex-specific manner. We denote male and female fecundity by *w*_m_(*z*) and *w*_f_(*z*) respectively. Unless otherwise stated, we assume that these are given by Gaussian functions: 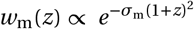 and 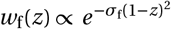 (as in e.g. Connallon and Clark, 2014a,b; Fisher, 1958; Lande, 1980; Muralidhar and Coop, 2024), such that male fitness is maximised by trait value *z =* −1 and female fitness by *z =* 1, where *σ*_m_ and *σ*_f_ determine how steeply fitness declines with deviation from the optimum in males and females respectively (i.e. *σ*_m_ and *σ*_f_ modulate the strength of selection in males and females).

### 2.2 A sexually antagonistic locus on the PAR

As a baseline, we assume *z* is influenced by a locus (“trait locus” or “z-locus”) in the pseudo-autosomal region (PAR) of a sex chromosome that recombines with a male SDR with probability *r* > 0 per meiosis. We therefore initially consider an XY system, which under the random union of gametes model of mating is equivalent to a ZW system with the sex labels exchanged. This symmetry breaks down when male and female reproductive variances differ owing to the mating system; in that case, we model ZW explicitly (see section 3.4).

Alleles at the trait locus have additive effects on phenotype, such that an individual’s trait value is given by *z = x*_1_ *+ x*_2_ where *x*_1_ is the allelic value carried on its maternally inherited X-chromosome and *x*_2_ is the allelic value carried on its paternally inherited chromosome (which is an X-chromosome in females and a Y-chromosome in males). Because alleles have the same phenotypic effect in both sexes but male and female fitness are maximised at different trait values (−1 in males and 1 in females), an allele that increases fitness in one sex can decrease fitness in the other; i.e. the trait locus is sexually antagonistic.

### 2.3 Invasion of a cis-recombination suppressor

We investigate the fate of a new mutation arising within the PAR that fully suppresses recombination between the SDR and the *z*-locus (such as an inversion capturing both regions) when the trait locus is polymorphic for sexually antagonistic alleles. We consider two cases of sexually antagonistic polymorphism.

1. We assume that two alleles, A and a, segregate at the trait locus with arbitrary effects *x*_A_ and *x*_a_, respectively, and that these effects maintain a balanced polymorphism at the trait locus; otherwise polymorphism would be transient and unlikely to be captured by a recombination suppressor. Without loss of generality, we assume *x*_A_ < *x*_a_, so that the A-allele tends to be male-beneficial and a-allele female-beneficial.
2. We focus on the situation where *x*_A_ and *x*_a_ have converged to a polymorphic evolutionary equilibrium via Kimura’s continuum-of-alleles model (Kimura, 1965); i.e. each mutation adds a small random deviation with mean zero to the allelic value it occurs on. In this model, sexually antagonistic selection can drive the gradual emergence of polymorphism via evolutionary branching (Appendix A.1; see Flintham et al., 2025 for more details about evolutionary branching on the PAR; also Siljestam et al., 2024; Van Dooren et al., 2004 for other genomic locations). This eventually results in the stable coexistence of two diverged allelic lineages whose equilibrium values we denote by 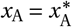 and 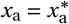 (Appendix A.2 explains how to characterise this polymorphism).

These two cases may reflect different timescales of evolution at the trait locus and the recombination modifier. Model (i) applies when mutations suppressing recombination and those influencing *z* arise on similar timescales. Model (ii) applies when mutations affecting allelic effects at the trait locus occur more frequently than mutations at recombination modifiers, so that the trait locus typically equilibrates before a new suppressor arises. Model (ii) may also capture a polygenic situation in which *z* is encoded by multiple PAR loci with fixed additive effects; we analyse such a model in Appendix B, where we obtain results analogous to those under the continuum-of-alleles model.

We initially assume the recombination suppressor appears on the Y-chromosome (i.e. is Y-linked) and captures the SDR itself. Later in section 3.3 we consider a suppressor appearing on the X-chromosome (i.e. X-linked) and capturing the homologous genomic region to the SDR on the X (equivalent modifiers in ZW systems are modelled in section 3.4).

To investigate the evolution of recombination arrest, we first calculate the invasion fitness 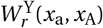 of a rare Y-linked suppressor in a resident population where recombination otherwise occurs at rate *r* between the SDR and the trait locus, and where alleles with effect sizes *x*_a_ and *x*_A_ segregate at their equilibrium frequencies (Appendix C.1 characterises 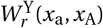, which formally is defined as the long-term geometric mean of the average per-capita number of suppressor copies produced per demographic time step). When 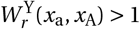, the suppressor can invade the population; otherwise, it goes extinct with probability one.

## 3 Results

### 3.1 Sexually antagonistic polymorphism on the PAR generates male heterozygote advantage

As a first step, we characterise the polymorphism that emerges through gradual evolution (continuum-of-alleles model; case (ii) in section 2.3) at the trait locus whilst keeping recombination rate fixed at *r*. Using a numerical approach (Appendix A.2, in particular Appendix A.2.2) we find that for all parameters considered, the evolutionary equilibrium always consists of two alleles – a and A – that respectively encode trait values 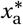 and 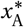 such that: (i) in females, aa homozygotes show the greatest fitness; (ii) in males, Aa heterozygotes show the greatest fitness (Fig. 1A-B for an example, and see Fig. 1C and SI Fig. 1 for 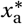 and 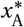 across a broad range of parameter sets). This occurs because one allele, a here, leads homozygotes to express a trait value close to the female optimum, while the other allele A leads homozygotes to express a trait value further from the male optimum than the one expressed by Aa heterozygotes (which occurs whenever *x*_A_ < −(2 *+ x*_a_)/3).

**Figure 1.**
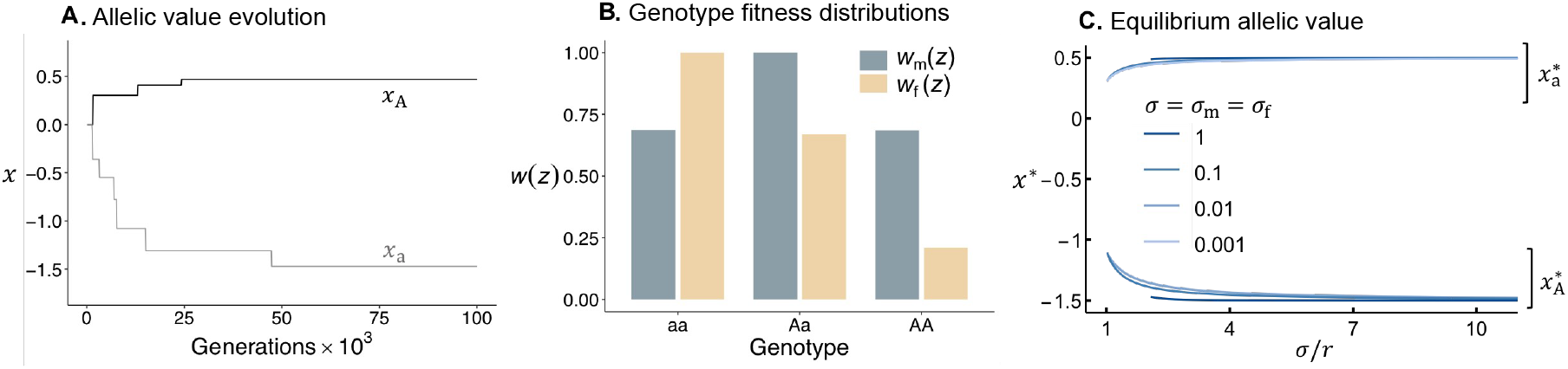
Evolution of allelic values at a sexually antagonistic locus on the PAR. Panel **A.** shows the evolution of the allelic values *x*_a_ and *x*_A_ from individual-based simulations (Appendix A for details). Plot shows the allelic values encoded by the two most frequent alleles at a given generation in a simulation replicate (which correspond to alleles a and A). These values diverge through the accumulation of recurrent mutations, until they converge at 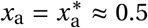 and 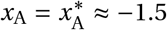. Other parameters used in simulation: *N =* 10^4^, *σ = σ*_m_ *= σ*_f_ *=* 0.1, *r =* 0.005, effects of new mutations are sample from a normal distribution with mean 0 and standard deviation 0.25. Panel **B**. shows the fitness (*w*_m_(*z*) and *w*_f_(*z*)) of different genotypes (aa, Aa and AA) at generation 10^5^ of the simulation in panel **A**, i.e., once allelic values have converged to their evolutionary stable values 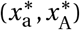. These show male heterozygote advantage. Panel **C**. shows the evolutionary stable values 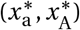 (see Appendix A for calculation) as a function of the ratio between the strength of sexually antagonistic selection and the recombination rate (*σ*/*r* where *σ = σ*_m_ *= σ*_f_; recall *r* < *σ* is the condition for disruptive selection in the PAR, hence why some curves do not cover the entire x-axis). Different curves show this for different strengths of *σ*. Regardless of the strength of sexually antagonistic selection, allelic values at the stable values are 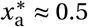 and 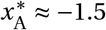 unless *σ*/*r* is close to 1 (which corresponds to where disruptive selection at the trait locus is weakest). Similar results are found when *σ*_m_ ≠ *σ*_f_ (see SI Fig. 1).

The “overshooting” of the male trait optimum by allele A occurs because once both alleles have begun diverging, Aa is the most frequent male genotype and hence selection on A favours these heterozygotes to be best adapted, at the expense of AA males. To see why, consider that once two alleles have diverged in their phenotypic effects, selection leads male-beneficial allele A to become over-represented on the Y-chromosome and hence in Y-bearing sperm, while female-beneficial allele a becomes overrepresented in X-bearing eggs. This increases the number of female (XX) offspring that are aa homozygotes, and an increase the number of male (XY) offspring that are Aa heterozygotes (meanwhile, the number of offspring that are AA homozygotes decreases, owing to the low frequency of A in eggs). Consequently, selection favours an allelic value *x*_A_ for A that is greater than the male optimum in homozygotes because this *x*_A_ confers high male fitness in heterozygotes when combined with an allelic value *x*_a_ that confers high female fitness in homozygous form (e.g., with *σ*_m_ *= σ*_f_, the combination of allelic values *x*_A_ *=* −1.5 and *x*_a_ *=* 0.5 on average encode the fittest phenotypes as they satisfy *x*_A_ *+ x*_a_ *=* −1, the male optimum, and *x*_a_ *+ x*_a_ *=* 1, the female optimum). Overdominance here is restricted to males (Gavrilets and Rice, 2006, for a model assuming this on autosomes). This differs from the sex-averaged heterozygote advantage generated by dominance reversal for fitness (where homozygotes show highest within-sex fitness but deleterious alleles are recessive in each sex; Connallon and Chenoweth, 2019; Fry, 2010), which is required to maintain balanced sexually antagonistic polymorphism on autosomes under weak selection (Kidwell et al., 1977).

While the evolution of male overdominance in the continuum-of-alleles model arises under the gradual evolution of allelic values through recurrent mutation, we also show in Appendix B that a similar pattern emerges when *z* is polygenic (encoded by multiple additive loci in the PAR) so long as recombination amongst loci is not too strong. In this case, selection quickly sorts standing variation to generate two predominant haplotypes: one linked to the X-chromosome, with allelic effects summing to half the female optimum, and another linked to the Y-chromosome, with effects overshooting the male optimum (in other words, X- and Y-linked haplotype values converge respectively to 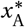 and 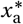). Female homozygote advantage (favouring two copies of the X-linked haplotype) and male heterozygote advantage (favouring an X-Y haplotype combination) thus ensues.

### 3.2 Sexual antagonism expedites sex chromosome evolution, but its potency is sensitive to the mating system

Next, we characterise the fate of a recombination modifier on the Y-chromosome that arises as a single copy and fully suppresses recombination between the SDR and the trait locus; which is polymorphic for alleles A and a. In Appendix C.1-C.2 we show that across combinations of allelic effects satisfying *x*_a_ > *x*_A_, the invasion fitness of such a suppressor always exceeds unity 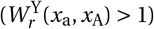 if it captures allele A and is less than unity 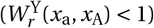 if it captures allele a. That is, selection favours recombination suppressors capturing A and disfavours suppressors capturing a (as previously shown for small-effect recombination modifiers; see the condition for selection to favour recombination suppression in Otto, 2014 with *R*_*m*_ *=* 0, and consistent with results from full suppressor models that constrain allelic values such that *x*_A_ is strictly male-beneficial, e.g., Charlesworth and Charlesworth, 1980; Nei, 1969; Olito and Abbott, 2025). In addition, we find that selection for recombination arrest is greatest (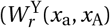 largest) when the resident recombination rate (*r*) is largest (see also Olito and Abbott, 2025) and is especially strong when alleles have evolved to their evolutionary equilibrium (i.e., when 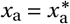 and 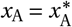, see Appendix C.2 for discussion).

The tempo of recombination evolution, however, depends not just on whether selection can favour a rare suppressor, but also on the likelihood that such a mutant evades stochastic loss and goes on to fix in the population. To quantify this, we first calculate the (unconditional) probability of invasion Ψ^Y^ of a Y-linked suppressor: the probability that a suppressor arises in a chromosome carrying allele A and that this suppressor does not go extinct in the long run, i.e. 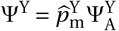, where 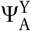 is the invasion probability of the suppressor conditional on it capturing A and 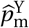 is the equilibrium frequency of A in sperm (see Appendix C.3.1 for details, and also Olito and Abbott, 2025 for a similar approach). Across all parameters tested, suppressors that do not go extinct (i.e, successfully invade) always go on to fix in the population; in other words, suppressors do not form balanced polymorphisms (Appendix C.3.2 for details). The probability of invasion, Ψ^Y^, is therefore equivalent to the probability of fixation, and this allows us to leverage Ψ^Y^ to compute a suppressor’s expected time to substitution, *τ* (the expected waiting time to arise and then fix in a population, Appendix C.3.2 for details). We use *τ* as a measure of the rate of recombination evolution.

We first perform this analysis under the life cycle described in section 2.1, where mating is modelled as a random-union-of-gametes model (such that the offspring distribution of a rare recombination suppressor is Poisson with mean given by eq. C-4). We calculated Ψ^Y^ for different allelic values *x*_a_ and *x*_A_, and parameters *σ*_m_, *σ*_f_ and *r* (Appendix C.3.1 for details). As expected from the results for the magnitude of 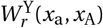 (Appendix C.2), invasion probability Ψ^Y^ is especially large when allelic values are at their evolutionary equilibrium (i.e., when 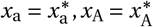, SI Fig. 4 and Fig. 3A compare dotted and solid lines), and greatest when sexual antagonism is strong and the resident recombination rate is high (*σ = σ*_m_ *= σ*_f_ and *r* large, Fig. 3A).

More generally, unless allelic values are very weakly diverged (*x*_a_ and *x*_A_ similar) or sexually antagonistic selection is very weak (*σ* very small), Ψ^Y^ is high compared to the neutral expectation in a large population (as also suggested by Olito and Abbott, 2025, by comparing the scale of Y-axes in their Figs. 2A-C). For example, with allelic values *x*_a_ *=* 0.5 and *x*_A_ *=* −0.5 and loose linkage *r =* 0.5, Ψ^Y^ ≈ 0.1 for *σ =* 0.1 and Ψ^Y^ ≈ 0.01 for *σ =* 0.01 (Fig. 3A). A neutral suppressor would fix with a similar probability only in very small populations (with 10 and 100 males or less for *σ =* 0.1 and *σ =* 0.01, respectively). In line with this, the expected time to substitution for a recombination suppressor is orders of magnitude shorter if it captures a sexually antagonistic locus than for a neutral suppressor (e.g., with reasonable suppressor mutation rates and *r =* 0.5, a substitution can be expected to occur roughly 1000 times faster than at neutrality when *σ =* 0.1, and around 100 times faster when *σ =* 0.01, Fig. 3B).

**Figure 2.**
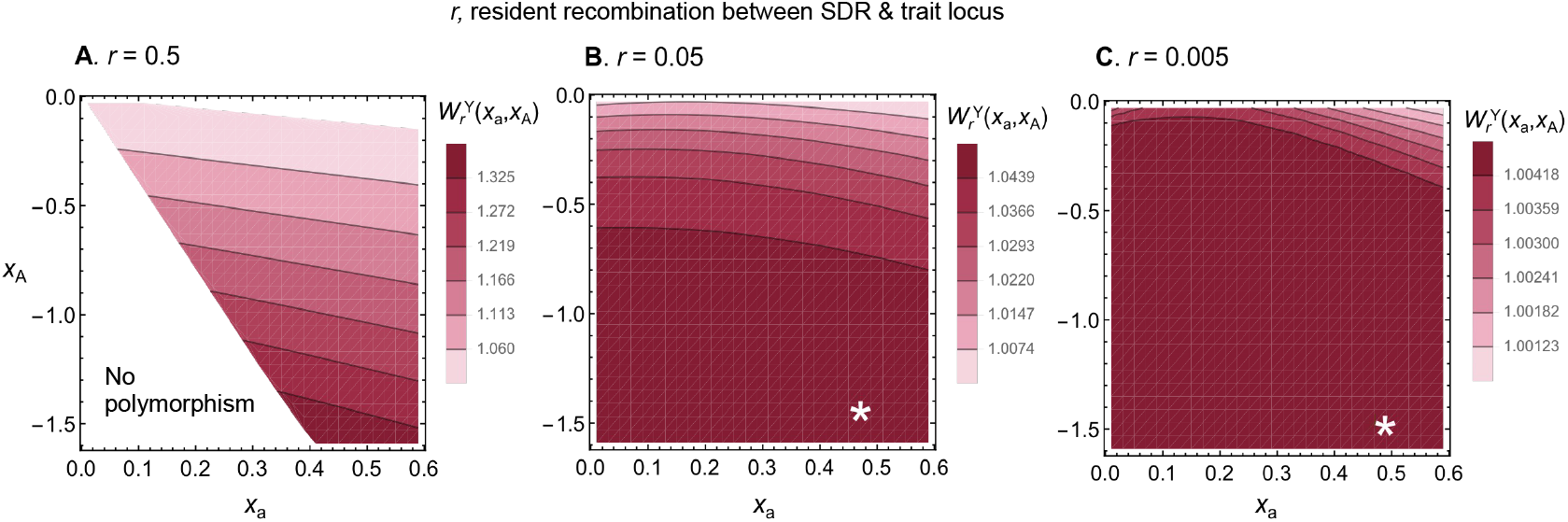
Invasion fitness of a Y-linked suppressor capturing a trait locus with alleles encoding *x*_a_ and *x*_A_. Plots show the invasion fitness, 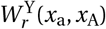 (main text eq. C-4, Appendix C.1 for calculation details), of a Y-linked recombination suppressor capturing allele A at the trait locus as a function of the allelic values *x*_a_ and *x*_A_. Different panels show results for different levels of resident recombination between the trait locus and the SDR, *r*. White stars show the location of the evolutionary stable allelic values that would emerge through gradual evolution 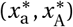 (there is no star in Panel **A**, because sexual antagonism does not produce disruptive selection here: *r* > *σ*, with balancing selection thus requiring large-effect mutations; see eq. A-2). Non-coloured space indicates that a suppressor cannot invade (which is because these conditions do not lead to a balanced polymorphism for alleles A and a at the trait locus). Other parameters used *σ = σ*_m_ *= σ*_f_ *=* 0.1 (see SI Fig. 3 for cases where *σ*_m_ ≠ *σ*_f_, and SI Fig. 2 for the selection and dominance coefficients of alleles A and a in the notation of Kidwell et al. (1977)).

The random-union-of-gametes model is a good approximation for large outbred populations where individuals of both sexes mate many times, but not for the many species experiencing sex-specific reproductive variance or skew (Andersson, 1994; Clutton-Brock and Parker, 1992; Shuker and Simmons, 2014). To capture such a situation, we now consider a lifecycle that begets a lottery polygyny mating system (Lee et al., 2008; Nunney, 1993). Briefly, we assume females mate singly but males can mate multiply (full details of this life cycle are given in Appendix C.3.3). A male’s likelihood of mating with each female is determined by his relative mating competitiveness, which is a random value assigned to him upon reaching adulthood (e.g. capturing his biological “condition”, Wilson and Nussey, 2010, which we assume is independent of genotype here, to facilitate comparison between models). Male reproductive variance therefore emerges because on average some males mate with more females, and this effect can be tuned by modifying the distribution of male mating competitiveness, with variance in mating competitiveness modulating the distribution of male mating and reproductive success (Lee et al., 2008). We detail how we calculate the probability of invasion Ψ^Y^ of a Y-linked suppressor under this model in Appendix C.3.3.

Comparison between Ψ^Y^ calculated under this model and under the random-union-of-gametes reveals that Y-linked recombination suppressors are less likely to invade under lottery polygyny (Fig. 3C). This is because high variability in male mating success elevates the probability that a male by chance mates no females and so produces no offspring. This in turn exacerbates the probability that the lineage carrying a recombination suppressor suffers stochastic extinction. Consequently, adaptive (i.e., those capturing allele A) suppressors will fix with probabilities similar to neutral suppressors in much larger populations under lottery polygyny than under the random-of-union-of-gametes model – especially when males show large variability in their mating success. For example, where male mating variance is around 10 (which is well within empirical estimates; Wade and Shuster, 2004 Table 1), population sizes of 10^4^ males will yield similar neutral and adaptive fixation probabilities when *r =* 0.5 and *σ =* 0.01. Accordingly, adaptive recombination suppression evolves more slowly in the presence of high male reproductive variance (Fig. 3D), and even on the same timescale as neutral suppression if sexual antagonism is sufficiently weak (Fig. 3D, *σ =* 0.001).

**Figure 3.**
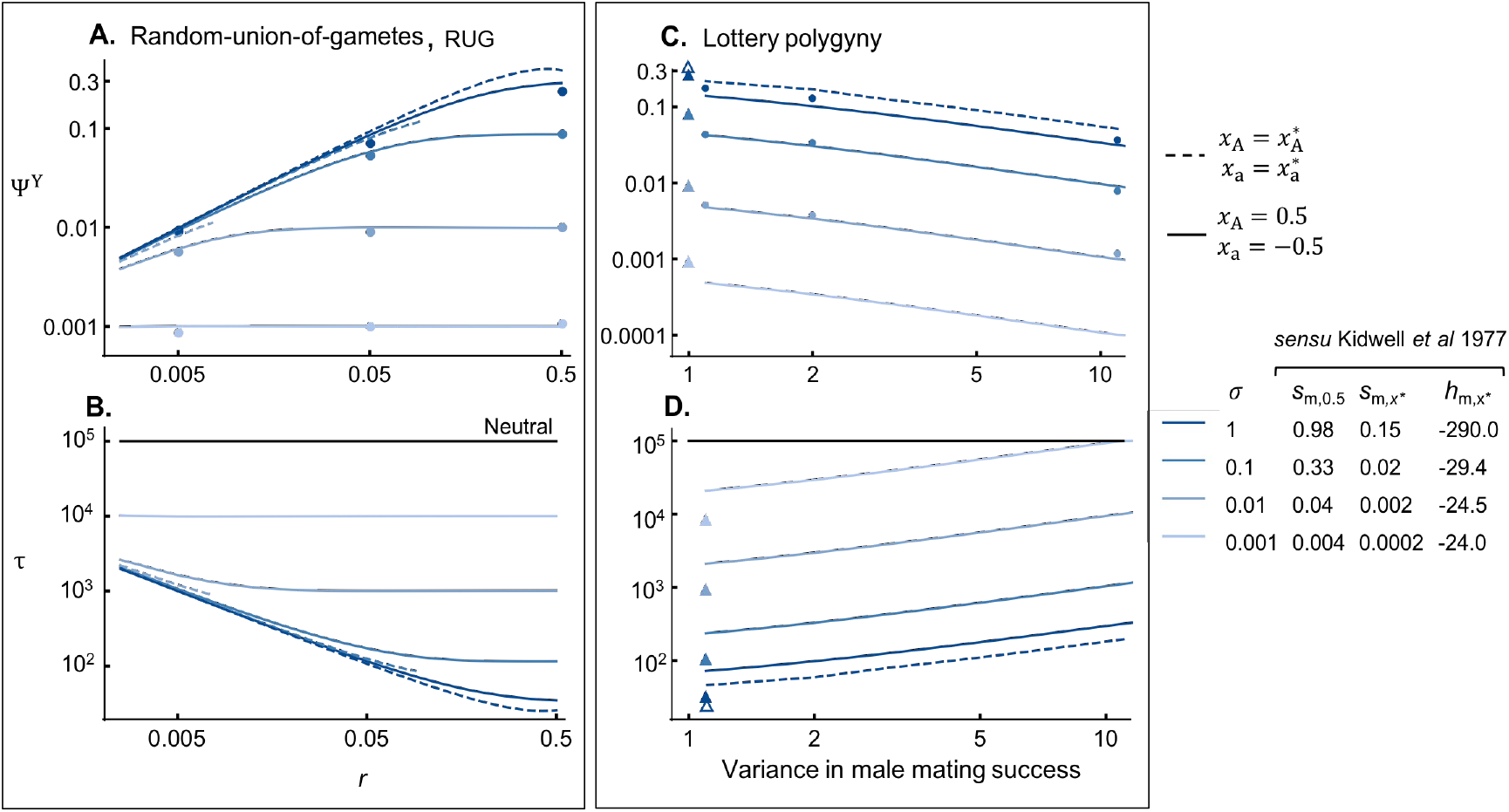
Invasion probability and waiting times of Y-linked suppressors. Panel **A.** shows the unconditional invasion probability of a newly arisen Y-linked suppressor, Ψ^Y^, under the random-union-of-gametes (RUG) model (Appendix C.3.1 for calculation) according to the resident recombination rate (*r*). Dashed lines correspond to invasion probabilities when allelic values have converged to their evolutionary stable values 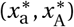 and are therefore shown only for the range of parameters where evolutionary branching occurs. Solid lines correspond to invasion probabilities when allelic values have diverged, but not so much that males show heterozygote advantage: *x*_a_ *=* 0.5 and *x*_A_ *=* −0.5. Dots show results from individual-based simulations with *x*_a_ *=* 0.5 and *x*_A_ *=* −0.5 (which correspond to proportion of replicate simulations in which a Y-suppressor fixed, Appendix C.3.2). Different colours correspond to different strengths of selection at the trait locus (*σ = σ*_m_ *= σ*_f_). Other measures of the strength of selection at the trait locus (following the convention of Kidwell et al., 1977) are also given in the figure table: (1) the male homozygous selection coefficient, *s*_m_ *=* [*w*_m_(2*x*_A_) − *w*_m_(2*x*_a_)]/*w*_m_(2*x*_A_), for allele a (where *s*_m,0.5_ refers to the case *x*_a_ *=* 0.5, *x*_A_ *=* −0.5, and 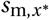 to the case 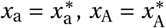); (2) the male dominance coefficient for allele a when 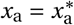 and 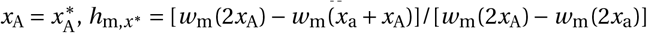 – negative values for *h*_m_ indicate male fitness overdominance, and we see large absolute values here because homozygotes show similar fitness when alleles are at their evolutionary stable values; i.e., 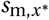 is very small. Panel **B**. shows the expected waiting time *τ* for a Y-linked suppressor to substitute into a population of 10^4^ males (i.e., *N =* 2*×*10^4^) with mutation rate *µ =* 10^−5^ (these are equivalent parameters to those considered by Lenormand and Roze, 2022, their Fig. 2, see Appendix C.3.2 for calculation and further discussion of mutation rates). The same parameter values are used as in panel **A**.. Horizontal black line shows the expected waiting time for a purely neutral Y-linked allele (1/*µ*). Panels **C**.-**D**. show the same output as **A.-B**. but for a model of lottery polygyny. In these panels we assume loose linkage *r =* 0.5 between the SDR and trait locus, and vary the degree of variance in male mating success (Appendix C.3.3 for calculation; note that evolutionary branching only occurs when *σ* > 0.5 here). Triangles show equivalent Ψ^Y^ and *τ* values under the random-union-of-gametes model (empty and filled triangles for 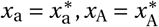 and *x*_a_ *=* 0.5, *x*_A_ *=* −0.5 cases, respectively).

**Table 1.** Predictions for patterns of sex chromosome evolution under different drivers recombination suppression. Predictions for the rate of recombination suppression when suppressors (i) capture a sexually antagonistic (SA) locus, (ii) capture sexually antagonistic (SA) and deleterious loci (iii) are neutral (iv) capture deleterious loci, under scenarios in which male and female reproductive variance is either equal (as in the random-union-of-gametes model) or male-biased (as in our lottery polygyny model). Columns are ordered according to the overall pace of sex chromosome evolution predicted under each driver (pure sexual antagonism being fastest and pure deleterious variation slowest). In this table X vs Y and Z vs W denote the relative substitution rates of the X and Z chromosomes, respectively, while XY vx ZW refers to the overall pace of recombination evolution in XY systems relative to ZW systems (by slight abuse of notation, > and < denote faster and slower recombination evolution, respectively, and ≳ denotes faster or similar rates). Blue highlighted cells are those with unique patterns for suppression driven by sexually antagonistic selection. All these baseline expectations are “all else equal”, i.e., assuming no differences in mutation rates, intrinsic costs of suppression, or strengths of selection between different mating systems, sex chromosomes, or sex-determination systems.

| Mating system | SA | SA & deleterious variation | Neutral | Deleterious variation |
| --- | --- | --- | --- | --- |
| Equal variance | $X \gtrsim Y$ | $X \gtrsim Y$ | $X = Y$ | $X \gtrsim Y$ |
| | $Z \gtrsim W$ | $Z \gtrsim W$ | $Z = W$ | $Z \gtrsim W$ |
| | $XY = ZW$ | $XY = ZW$ | $XY = ZW$ | $XY = ZW$ |
| Male-biased variance | $X > Y$ | $X > Y$ | $X = Y$ | $X \approx Y$ |
| | $Z < W$ | $Z < W$ | $Z = W$ | $Z \approx W$ |
| | $XY < ZW$ | $XY < ZW$ | $XY = ZW$ | $XY = ZW$ |

### 3.3 Sex chromosome evolution is more readily driven by X-than Y-linked suppressors under sexual antagonism

We next investigate the potential for recombination evolution to occur through X-versus Y-linked modifiers. Divergence between sex chromosomes may just as well be mediated by recombination modifiers fixing on the X as the Y (e.g., Charlesworth and Charlesworth, 2005; Lenormand and Roze, 2022; Olito and Abbott, 2025, see also Lenormand, 2003; Otto, 2014 for gradual suppression models that are agnostic as to the genomic location of modifiers), but some authors have suggested that X-driven suppression is unlikely because X-linked suppressors will be more weakly selected than those on the Y (see below, and Blackmon and Brandvain, 2017; Charlesworth and Charlesworth, 1980; Olito and Abbott, 2025, for modelling support). To study recombination evolution on the X-chromosome, we now consider the invasion of a suppressor on the X-chromosome across different allelic values (*x*_a_ and *x*_A_, including their evolutionary equilibrium 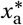 and 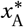) and mating systems (random-union-of-gametes and polygyny models). We assume this modifier fully suppresses recombination in both males and heterozygous females (as for a chromosomal inversion, Kirkpatrick, 2010).

We first calculate the invasion fitness of an X-linked suppressor, 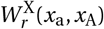 (in Appendix D.1). Analysis of 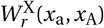 shows that such a suppressor is favoured (i.e., 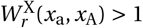) for any parameters *σ*_m_, *σ*_f_ and *r*, and allelic values *x*_a_ and *x*_A_ such that a and A are maintained under balancing selection, if and only if the suppressor captures female-beneficial allele a. This is because linking allele a to the SDR-homologue on the X-chromosome ensures that males always transmit this allele to their female offspring, who then enjoy higher fitness as a result. As with Y-linked suppression, selection favours an X-linked suppressor most strongly (i.e., 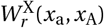 is largest) when recombination in males is frequent (*r* large), with selection especially strong when alleles are at their equilibrium values (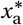 and 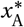) (SI Fig. 5).

Selection therefore readily favours X-linked suppression, but we find it does so less strongly than for suppression on the Y (i.e., 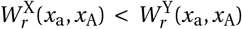, SI Fig. 6A, consistent with the results of Charlesworth and Charlesworth, 1980 and Blackmon and Brandvain, 2017). The selective advantage of an X-linked suppressor is approximately three times weaker than the advantage of a Y-linked suppressor when selection for recombination suppression weak (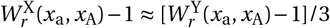 when *σ*_f_ and *σ*_m_ small or *r* small). This is primarily because the X spends 2/3 of its time in females where recombination suppression is inconsequential to offspring fitness (as chromosome segregation in females is random with respect to offspring sex, Blackmon and Brandvain, 2017; Charlesworth and Charlesworth, 1980). This dilutes the fitness advantage of X-linked recombination suppressors relative to those on the Y, which are passed strictly through males. An additional penalty to X-linked suppression arises for non-equilibrium allelic values where males do not show heterozygote advantage and thus ubiquitously favour allele A, as here an X-suppressor capturing allele a will confer decreased fitness when carried in males.

Next we calculate the probability of invasion Ψ^X^ for an X-linked suppressor of recombination and compare it with its Y-linked counterpart Ψ^Y^ (Appendices D.2.1-D.3 for full description of invasion probabilities for X-linked suppressors; note that some of the following results contradict those in Appendix C of Olito and Abbott, 2025, which is due to a technical error in Olito and Abbott, 2025 that is corrected here, see Appendix D.2.2 for details). Where the offspring distribution of a rare X-linked suppressor is Poisson (as expected under the random-union-of-gametes model), we find that such a suppressor is less likely to invade than a suppressor arising on the Y, i.e. Ψ^X^ < Ψ^Y^ (SI Fig. 6B). This is unsurprising given that selection favouring X-linked suppression is weaker (i.e., 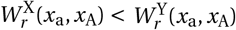).

Nevertheless the discrepancy between X and Y invasion probabilities remains moderate – equal to or less than three-fold (SI Fig. 6B) – and can therefore be more-than compensated for by stronger mutational input of X-linked inversions: when recombination suppressors arise with equal per-base rates in males and females, new suppressors are three times more likely to arise on the X than Y owing to the X’s larger population size. Therefore, the probability that a recombination suppression event (i.e., substitution by a suppressor) is due to a Y-rather than an X-linked modifier is *ρ*_YX_ *=* Ψ^Y^/(Ψ^Y^ *+* 3Ψ^X^) (provided suppressors arise rarely). Plugging in our values for Ψ^Y^ and Ψ^X^ into *ρ*_YX_ reveals that recombination suppression is in fact equally likely to be X- or Y-linked when sexual antagonism is weak or initial linkage between the SDR and trait locus is tight (i.e., *ρ*_YX_ *=* 0.5 when *r* or *σ* small, Fig. 4A), because the three-fold higher selective advantage to Y-linked suppressors cancels with the discrepancy in mutational input. Otherwise, we find X-linked suppression is more probable (e.g., *ρ*_YX_ < 0.5 with *r =* 0.5 and *σ* ≥ 0.1 for *x*_a_ *=* 0.5, *x*_A_ *=* −0.5 and 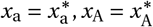, Fig. 4A). This is because when sexual antagonism and prior recombination are strong (*σ* and *r* large), the selective advantage to Y-linked suppression is constrained by the combined effects of (i) beneficial alleles tending to be more recessive under Gaussian fecundity, and (ii) the excess of suppressor-carrying males who are Aa heterozygotes (relative to X-linked suppressor-carrying female aa homozygotes, see section 3.1), who therefore experience a weaker benefit of guaranteed paternal inheritance of allele A. Y-linked suppressors thus show a below-three-fold selective advantage relative to X-linked suppressors and so a lower substitution rate (*ρ*_YX_ < 0.5).

**Figure 4.**
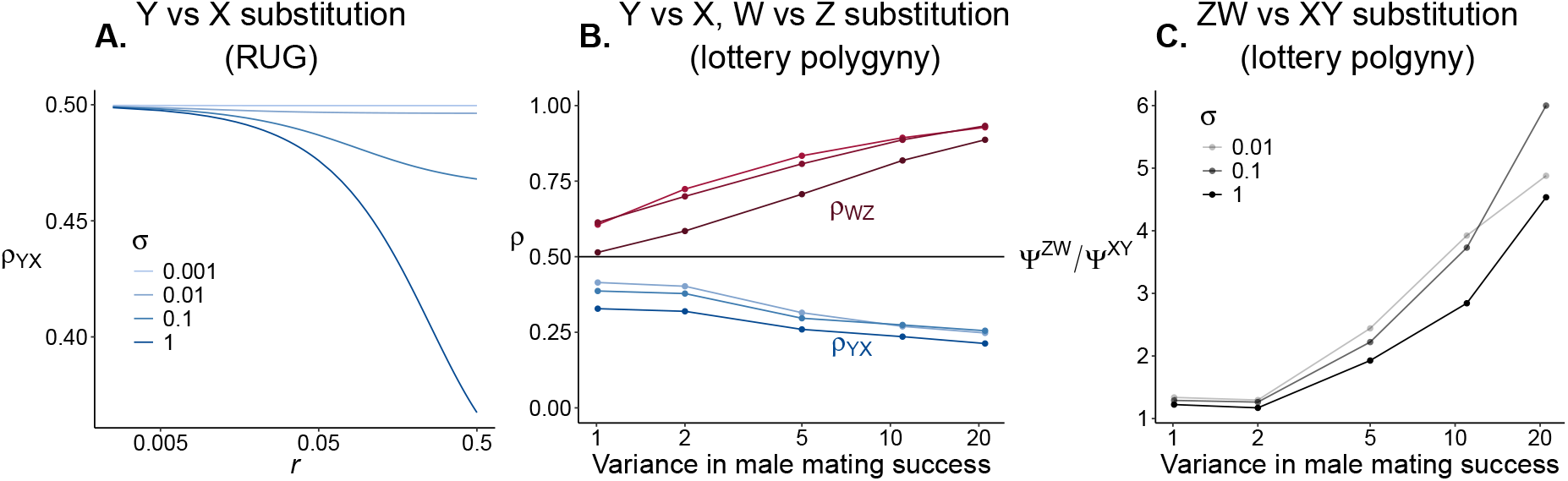
Likelihood of heterogametic-vs. homogametic-linked suppression and pace of evolution in XY vs. ZW systems. Plots show the probability that a substitution by a recombination suppressor is on the heterogametic versus homogametic chromosome (Y-vs. X-linked: *ρ*_YX_ *=* Ψ^Y^/(Ψ^Y^ *+* 3Ψ^X^) or W-vs. Z-linked: *ρ*_WZ_ *=* Ψ^W^/(Ψ^W^ *+* 3Ψ^Z^), Appendices C.3.1 and D.2.1 for calculation of Ψ^X^ and Ψ^Y^ in the random-union-gametes model, respectively). Panel **A.** shows *ρ*_YX_ under the random-union-of-gametes (RUG) model for different strengths of sexually antagonistic selection (*σ = σ*_m_ *= σ*_f_, see also Fig. 3 for other measures of the strength of selection) and resident recombination rate *r*. Lines show results for allelic values *x*_a_ *=* 0.5, *x*_A_ *=* −0.5 Note that in this model XY and ZW systems are interchangeable (*ρ*_YX_ *= ρ*_WZ_). Panel **B**. shows *ρ*_YX_ (blue curves and dots) and *ρ*_WZ_ (red curves and dots) output but for the polygyny model with loose linkage (*r =* 0.5) and different strengths of male mating variance. Here, fixation probabilities are estimated from individual-based-simulations (coloured dots are individual estimates). Horizontal line marks *ρ*_YX_ *=* 0.5. Panel **C**. shows ratio between the overall substitution rate in ZW systems relative to XY systems, Ψ^ZW^/Ψ^XY^ where Ψ^ZW^ *=* Ψ^W^ *+* 3Ψ^Z^ and Ψ^XY^ *=* Ψ^Y^ *+*3Ψ^X^, for different male mating variances. Dots are again estimated from individual-based simulations. See Fig. 3 for corresponding parameter values for selection at sexually antagonistic loci in Kidwell et al. (1977) notation.

Computing *ρ*_YX_ under lottery polygyny shows that recombination suppression is even more likely to be a consequence of X-linked suppressors under this mating system (Fig. 4B). In particular, when male reproductive variance is high, recombination arrest among sex chromosomes is expected to be mostly due to X-linked factors (e.g. with a variance in the number of mates among males of around 10, *ρ* ≈ 0.2, Fig. 4B). This is because male reproductive variance has less influence on the level of drift experienced by X-than Y-linked mutations so that, all else equal, X-linked suppressors are less susceptible to stochastic extinction than Y-linked suppressors here. By contrast, neutral substitution rates are always equal on the X and Y (as differences in mutational pressure and fixation probabilities balance out), leading to equivalent X-Y contributions to sex chromosome evolution through the fixation of neutral suppressors.

### 3.4 Sexual antagonism strongly promotes ZW recombination evolution through W-linked suppressors

In populations showing equivalent reproductive variance in males and females, our XY random-union-of-gametes model also exactly captures ZW sex-determination. Our results therefore predict equal, or mildly Z-biased, contributions to sex chromosome evolution in such populations (*ρ*_WZ_ *=* Ψ^W^/(Ψ^W^ *+* 3Ψ^Z^) ≈ 0.5 or *ρ*_WZ_ < 0.5). However, in species with sex-specific reproductive variance, evolutionary rates on different heterogametic (X vs Z) and homogametic (Y vs W) sex chromosomes can diverge. To investigate the consequences of this for sex chromosome evolution, we ran simulations of our polygyny model with a ZW system; i.e., where females carry a dominant SDR and males are homozygous for the SDR-homologue (App. D.3).

In ZW systems, elevated male reproductive variance leads to a very strong W-bias in the contribution to recombination arrest (Fig. 4B), owing to strong genetic drift through males dampening the efficacy of selection favouring Z-linked modifiers. Fixing suppressors are expected to almost always be W-linked when reproductive variance is high (e.g. with a variance in the number of mates among males of around 10, *ρ*_WZ_ ≈ 0.9, Fig. 4B). The degree of W-bias is greater than the degree of X-bias for the same mating system, because (i) suppressors on the W-chromosome are under stronger selection than those on the X, and (ii) unlike X-chromosomes, W-chromosomes are never present in males and so W-linked suppressors do not experience any exacerbation in drift due to polygyny at all.

Comparing the time to substitution for a recombination suppressor in an ZW system (which is inversely proportional to Ψ^ZW^ *=* 3Ψ^Z^ *+* Ψ^W^) to that in an XY system (inversely proportional to Ψ^XY^ *=* 3Ψ^X^ *+*Ψ^Y^), we see that in the presence of excess male reproductive variance, sexual antagonism drives faster recombination suppression in ZW systems (Ψ^ZW^ is greater than Ψ^XY^, Fig. 4C). This is again because the Y-chromosome is more strongly selected than the Z-chromosome (i.e., has larger fixation probability under random-union-of-gametes model; Ψ^Y^ > Ψ^X^ *=* Ψ^Z^), such that the effect of polygyny in eroding the efficacy of selection for modifiers on the Y disproportionately diminishes total substitution rates in XY systems relative to ZW.

### 3.5 Deleterious variation can hinder fixation on X and Z chromosomes

As a final step, we consider the joint effects of sexual antagonism and deleterious variation in the PAR on selection for recombination suppression. To do this, we ran multilocus simulations of a sex chromosome region comprising the SDR (or its homologue) and a section of the PAR containing a (i) single sexually antagonistic locus (with alleles A and a segregating) and (ii) a fixed number, *L*, of loci where deleterious alleles can appear (see Appendix E for full simulation details). Alleles at the sexually antagonistic and deleterious loci contribute multiplicatively to fitness, with each deleterious allele encoding a proportional reduction to fecundity of *s* and *hs* in homozygous and heterozygous form, respectively. Simulations are run until mutation-selection-drift balance at deleterious loci is reached before a recombination modifier is introduced, meanwhile, allele A is assumed to be at its equilibrium frequency. We then consider the fate of a suppressor that captures the entire chromosomal region with either the SDR (constituting a Y- or W-linked suppressor) or the SDR-homologue (constituting an X- or Z-linked suppressor) and again inhibits recombination when in heterozygous form, as for an inversion. We estimate fixation probabilities for each type of suppressor (X, Y, Z, W) across a range of parameters characterising deleterious (*s, h, L*) and sexually antagonistic (*σ*) variation.

In line with previous work (Charlesworth and Olito, 2024; Olito and Abbott, 2025; Olito et al., 2024; Roze and Lenormand, 2025), we find that in the absence of sexual antagonistic selection, deleterious variation generally generates below-neutral fixation probabilities for recombination modifiers on the Y- or W-chromosomes (Fig. 5A and SI Fig. 7A grey dots; above-neutral probabilities require deleterious alleles have weak effects and are highly recessive – *sh* is very small – leading to the “sheltering effect”, Jay et al., 2024; see Roze and Lenormand, 2025 for in-depth treatment). Furthermore, we also observe a pattern of below-neutral fixation of X- and Z-linked modifiers, although this reduction is less severe than for modifiers on the Y or W (SI Fig. 7B grey dots). This discrepancy is due to Muller’s ratchet acting on the backgrounds captured by “lucky” Y- and W-modifiers, which accelerates the accumulation of deleterious variants and hampers suppressor fixation (Roze and Lenormand, 2025). Muller’s ratchet is not seen in backgrounds captured by X- and Z-modifiers, as they still experience recombination in homozygotes.

**Figure 5.**
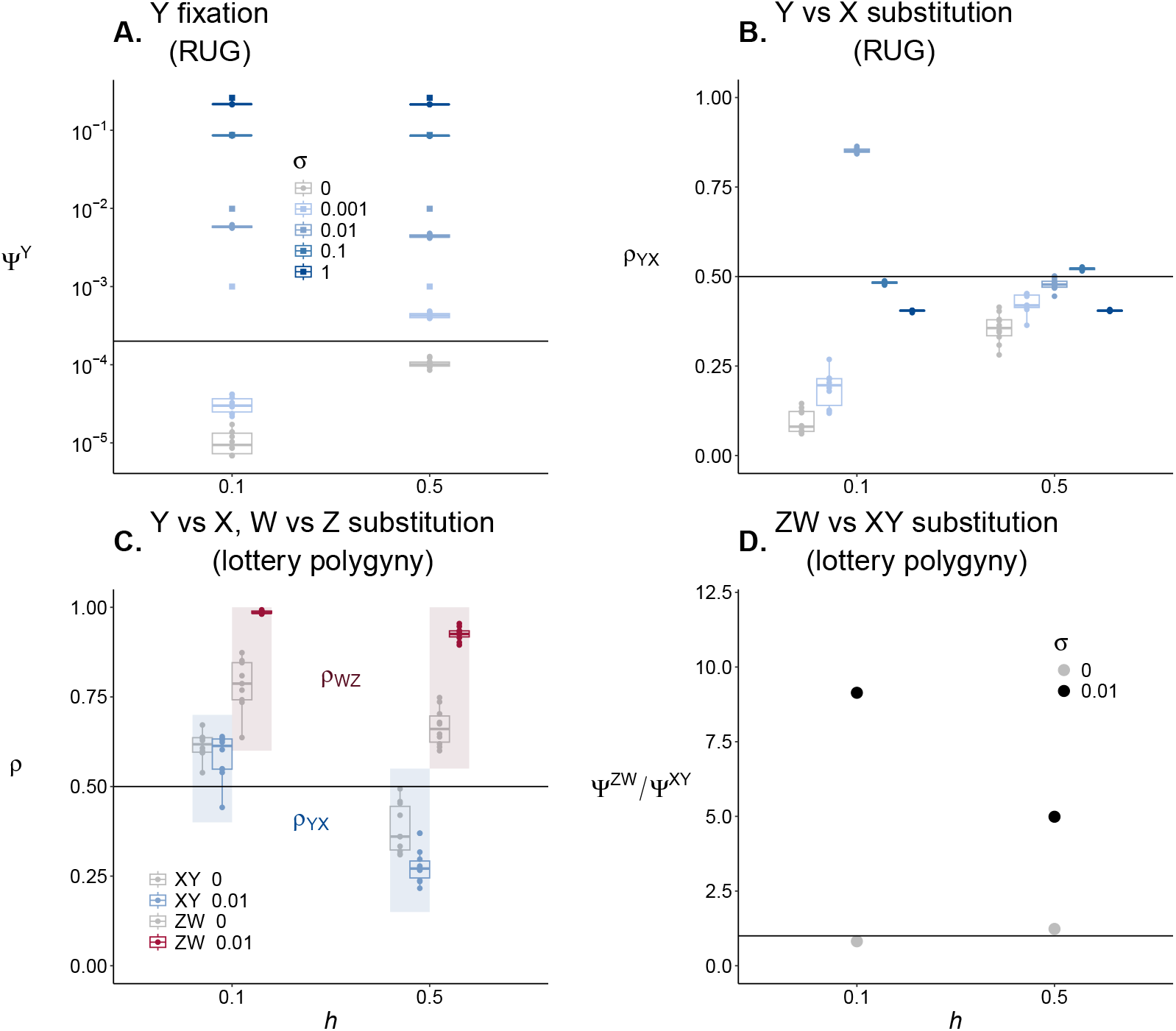
Selection for recombination suppression when deleterious and sexually antagonistic alleles co-segregate. Panel **A.** shows the fixation probabilities estimated from individual-based-simulations of a Y-linked suppressor capturing both a sexually antagonistic locus and *L =* 100 deleterious loci at mutation-selection balance (Appendix E for simulation details). Results are shown for selection coefficient *s =* 0.025 and dominance coefficients of either *h =* 0.1 and *h =* 0.5 at all deleterious loci (see x-axis). Each dot is an estimate for a single replicate population in a simulation of the random union-of-gametes model. Boxplot and dot colours correspond to different strengths of sexually antagonistic selection (*σ*). Squares show the invasion probability in the absence of deleterious variation (Fig. 3A). Horizontal line is the neutral fixation probability for a Y-linked mutation (1/(*N* /2)), where *N =* 10^4^. Recombination is scaled such that *r =* 0.3 between the sexually antagonistic locus and SDR (Appendix E). Panel **B**. shows the probability that a substitution by a recombination suppressor is Y-vs. X-linked, *ρ*_YX_, from the same simulation setup as in panel A (here XY and ZW systems are interchangeable: *ρ*_YX_ *= ρ*_WZ_). Horizontal line marks *ρ*_YX_ *=* 0.5. Panel **C**. shows the probability a substitution is on the heterogametic chromosome (*ρ*_YX_ or *ρ*_WZ_) in the lottery polygyny model with high male mating variance (variance ≈ 20) for different strengths of sexually antagonistic selection and dominance coefficients at the deleterious loci. Boxes and points in blue shaded regions show results for an XY system (show *ρ*_YX_) and boxes and points in red shaded region show results for a ZW system (show *ρ*_WZ_), grey and coloured boxes/points correspond to *σ =* 0 and *σ =* 0.01 cases, respectively. Panel **D**. shows overall substitution rates in XY and ZW systems with same parameters as in C, where each line is the average of all populations in the ZW simulations relative to the average of all populations in XY simulations (horizontal line marks Ψ^ZW^/Ψ^XY^ *=* 1). See Fig. 3 for corresponding parameter values for selection at sexually antagonistic loci in_27_Kidwell et al. (1977) notation.

When deleterious and sexually antagonistic alleles co-segregate, we find lower fixation probabilities relative to when sexually antagonistic alleles segregate in isolation (Fig. 5A). Deleterious variation hinders the fixation of suppressors here because the fitness benefit of picking up a beneficial sexually antagonistic allele (a for X- and Z-linked suppressors, and A for Y- and W-linked suppressors) can be masked if the suppressor is “unlucky”, i.e., it also captures a particularly loaded genetic background, or accumulates deleterious alleles during its segregation. Nevertheless, fixation probabilities for suppressors capturing a sexually antagonistic locus remain well above those for suppressors capturing deleterious loci alone, even when sexually antagonistic selection is weak. In other words, suppressors predominantly experience selection according to the allele they capture at the sexually antagonistic locus, because these loci account for a far greater proportion of the variance in fitness among genetic backgrounds than deleterious loci at mutation-selection-drift balance. Waiting times for the substitution of recombination suppressors will thus tend to be orders of magnitude shorter in the presence of sexual antagonism.

We also find that unbiased, X/Z-biased, and (more rarely) W/Y-biased substitution rates are all possible when deleterious and sexually antagonistic variation co-exist (Fig. 5B and SI Fig. 7C). In general, when male and female reproductive variances are equal, substitution rates are similar for modifiers arising on the hetero- and homogametic sex chromosomes if sexual antagonism is appreciable (*ρ* ≈ 0.5, in line with results presented in section 3.3). Meanwhile, X/Z-biased substitution is observed when sexual antagonism is very weak or absent, in which case fixation is driven by the capture of deleterious variation. Here, a combination of opposing effects: (i) Muller’s ratchet on the Y/W versus (ii) higher starting frequencies for Y/W- than X/Z-linked suppressors, lead to relatively similar fixation probabilities between hetero- and homogametic suppressors, and so overall to a X/Z-biased substitution rate (the exception being where *sh* is low enough to drive strongly Y/W-biased fixation rates through the sheltering effect, SI Fig. 7C with *s =* 0.005, *h =* 0.1).

Finally, our previous findings of divergence in genomic patterns between XY and ZW systems under male-biased reproductive variance also hold in the presence of deleterious variation. First, sexual antagonism can again lead to homogametic-biased (X-biased) substitution in XY systems, but heterogametic-biased (W-biased) substitution in ZW systems (Fig. 5C). Second, overall substitution rates in ZW systems are also larger than those in XY systems (Fig. 5D). These discrepancies between XY and ZW systems were either weak or absent when suppression evolved without sexual antagonism (Fig. 5C-D).

## 4 Discussion

### 4.1 Different timescales for sex chromosome evolution with versus without sexual antagonism

Our analyses indicate that recombination suppressors capturing sexually antagonistic loci generally have a substantially higher probability of invasion (and correspondingly lower expected time to substitution) compared with neutral suppressors or lucky suppressors capturing few deleterious alleles. Significant between-taxa variation exists in the rate at which recombination suppression evolves along sex chromosomes (Ma and Rovatsos, 2022). Our results thus suggest that systems evolving at different rates are being driven by different forces: fast suppression evolution associated with the segregation of sexually antagonistic alleles, while slow evolution proceeds due to the rare fixation of neutral or lucky suppressors. Another finding of our study is that the capture of a sexually antagonistic locus can appreciably influence the fixation probability of a recombination suppressor even when sex-specific selection is very weak and thus difficult to detect empirically. Because such loci may persist readily as balanced polymorphisms when they are located in the PAR, sexual antagonism can thus play a role in driving sex chromosome evolution even when its other effects are imperceptible.

Sexually antagonistic selection is particularly potent in driving recombination suppression when males show heterozygote advantage (for XY systems), which we have shown to be the evolutionary equilibrium for an antagonistic locus on the PAR. Alleles may reach this equilibrium through repeated small-effect mutations, as in the continuum-of-alleles model, though this process may be slow. But male heterozygote advantage could evolve much more rapidly, for instance, through large-effect mutations – in fact, mutations that overshoot the male optimum in a single step have the highest invasion fitness and are thus the most likely to invade – or through the evolution of standing variation in a polygenic trait encoded on the PAR (Appendix B). Male heterozygote advantage is therefore a plausible outcome of sexually antagonistic selection on the PAR, which in turn facilitates the evolution of recombination suppression.

### 4.2 Recombination arrest may be driven by suppressors on heterogametic or homogametic sex chromosomes

Our analyses also revealed a somewhat unexpected pattern: recombination suppression can more often be driven by the fixation of homogametic (X-linked or Z-linked) rather than heterogametic (Y-linked or W-linked) modifiers. Such a pattern arises when recombination suppression is driven by sexual antagonism in populations where both sexes show similar fitness distributions. This is observed despite suppressors appearing on the Y or W chromosome exhibiting a stronger selective advantage than those appearing on the X or Z (Blackmon and Brandvain, 2017; Charlesworth and Charlesworth, 1980). This is because the difference in positive selection for these two types of suppressor is insufficient to offset the three-fold discrepancy in their mutational input (owing to the homogametic chromosomes’ larger copy numbers).

This bias towards X-linked or Z-linked suppression is mild, but can be amplified for XY systems in taxa that show high male reproductive variance, because selection is less effective for Y-linked mutations due to the Y’s reduced effective population size. In ZW systems, however, elevated male reproductive variance instead produces the reverse pattern: a strong W-bias in suppression, which is because such variance exacerbates drift on the Z-chromosome and does not influence the W. A bias towards X- or Z-linked modifiers is also observed when recombination arrest is solely due to the fixation of lucky inversions. The precise level of X/Z bias here is the outcome of multiple factors, including Muller’s ratchet acting on Y and W inversions (Roze and Lenormand, 2025), and the larger initial frequency of heterogametic mutations. By contrast, when recombination suppression evolves neutrally (Ironside, 2010), evolutionary rates on the X and Y will be equivalent because X-Y differences in the fixation probabilities of neutral alleles exactly cancel differences in mutational input, irrespective of sex-specific drift.

By showing that suppressors on the homogametic sex chromosome may be at least as influential as those on the heterogametic chromosome in driving recombination suppression, our results provide a potential explanation for the empirical observations of X- and Z-linked inversions along sex chromosomes in a wide range of taxa (e.g., in fish, plants, and flies, Liu et al., 2025; McAllister, 2003; Moraga et al., 2025; Yue et al., 2023). In particular, Z-linked inversions are thought to have contributed significantly to the evolution of recombination suppression in birds (Zhou et al., 2014). While current data remains limited for drawing firm conclusions (Liu et al., 2025), comparing the distribution of sex chromosome inversions across homo- and hetero-gametic chromosomes, especially in species with well-characterised mating systems, may offer a promising approach to delineate the roles of selective and non-selective processes in driving sex chromosome evolution. According to our findings, if such comparisons were to reveal a bias towards X-linked inversions in XY systems and W-linked inversions in ZW systems across species with high male reproductive variance, this would implicate sexual antagonism as driving sex chromosome evolution.

Our results can also provide insight into the relative rates of sex chromosome evolution in XY and ZW systems. In species with stronger male reproductive variance, we show that adaptive recombination suppression evolves more rapidly in ZW than XY systems (all else being equal), as efficient selection in females relative to males leads to higher substitution rates for suppressors capturing female-beneficial alleles on the W-chromosome. Meanwhile, recombination suppression without sexual antagonism (i.e., driven by neutral or lucky inversions) should proceed equally quickly in XY and ZW systems. Because recombination arrest promotes genetic differentiation between the hetero- and homogametic sex chromosomes—often leading to W and Y degeneration (Bachtrog, 2013), this would suggest that heteromorphic sex chromosomes may evolve more rapidly in systems with ZW than XY sex-determination when driven by sexual antagonism. Moreover, because sex chromosome divergence is thought to stabilise sex-determination systems in the face of the appearance of new sex-determining loci (Pokorna and Kratochvíl, 2009; Vicoso, 2019) (although see Blaser et al., 2013, 2014 for a contrasting theory), our results would suggest that ZW systems evolving with sexual antagonism may also turnover less frequently than XY systems in polygynous species. Taken together, these results generate predictions for how rates of recombination suppression evolution should differ among X and Y, Z and W, and XY and ZW systems depending on mating system and on whether regions of recombination suppression is influenced by sexual antagonism, deleterious variation, or occurs neutrally; we summarise these patterns in Table 1.

### 4.3 Limitations and possible extensions

Our study considered explicitly cis-acting recombination modifiers (i.e., those that are in linkage to the SDR and trait loci) such as sex chromosome inversions (Kirkpatrick, 2010). One relevant factor of inversion biology that we did not consider is that these structural mutations can also induce fitness costs when present in heterozygous form, owing to the production of inviable gametes or zygotes (Berdan et al., 2023, for review). Importantly, if these costs act sex-specifically (which they may well do) then they could have a strong influence on the relative fitnesses of recombination suppressors on different sex chromosomes, and so on the balance of X(Z)-driven vs Y(W)-driven recombination evolution. For example, gamete inviability disproportionately harms females in many species (as sperm production is more weakly correlated with reproductive success than egg production unless males experience strong postcopulatory sexual selection, Janicke et al., 2016; Lehtonen, 2022), and so will penalise the invasion of X- or W-linked suppressors relative to those on the Y or Z. This effect could be sufficient to reverse the X- and W-bias in the basis of recombination suppression evolution. We therefore suggest that incorporating costs of suppression into recombination modifier models may be an important step towards understanding selection on inversions arising on different sex chromosomes.

In addition to modifiers acting in cis, recombination can be suppressed by trans-acting factors (Charlesworth, 2017; Jay et al., 2024) that are unlinked from the SDR and the trait locus (e.g., autosomal modifiers). In such cases, sex-specific selection can have diverse outcomes on recombination rates. In XY systems, selection on trans-acting modifiers can in fact favour the preservation, rather than suppression, of recombination on sex chromosomes. This occurs when males exhibit heterozygote advantage at a trait locus, but females show homozygote advantage for the allele that is at high frequency on the Y (Otto, 2014). However, we find that such a situation does not arise when allelic values diverge through gradual evolution, because the male-beneficial/female-deleterious allele is necessarily the one that increases in frequency on the Y (although it might arise if a large-effect female-beneficial mutation were introduced via male-biased introgression). Our results therefore suggest the evolution of allelic values in the PAR leads to polymorphism that favours recombination suppression by both cis- and trans-acting modifiers.

In this study, we have assumed that modifiers always suppress recombination, and we have focused on the rate at which such suppressors substitute in the population. It is possible, however, that mutations arise that undo recombination suppression once it has been established, namely chromosomal reversions. If (i) such reversions are sufficiently frequent and (ii) there are constraints on mechanisms that stabilise recombination suppression, such as dosage compensation (Lenormand and Roze, 2022, 2024), then reversions could slow sex chromosome evolution (Jay et al., 2024; Lenormand and Roze, 2024). Reversions would be especially consequential for neutral or lucky suppressors, because these modifiers are not maintained by selection once they have fixed. A reversion would therefore be neutral or even favoured if restoring recombination improves the efficacy of selection against deleterious mutations accumulated on the non-recombining background through Hill–Robertson interference (Lenormand and Roze, 2024). This effect should be strongest for reversions on the heterogametic sex chromosome, where recombination arrest is complete and interference is greatest. By contrast, the potential advantage of a reversion on the homogametic chromosome is less clear, because that chromosome continues to recombine and therefore experiences weaker interference. Instead, restoration of recombination could be favoured if, after an inversion has fixed on the X/Z, a perfectly homologous inversion subsequently arises on the Y/W, re-establishing collinearity between the sex chromosomes. More generally, adaptive suppressors capturing sexually antagonistic loci should be more robust to suppression-reversing mutations, although reversions may still invade if the accumulation of deleterious variation on the non-recombining background becomes sufficiently strong. While the prevalence of reversions on sex chromosomes remains an open question, extending previous work (Lenormand and Roze, 2022, 2024) to examine their effects on rates of recombination suppression in the presence of both sexual antagonism and deleterious variation would help clarify their potential role in sex chromosome evolution.

## 5 Conclusion

Sexually antagonistic selection has long been considered a key driver of sex chromosome evolution (Charlesworth and Charlesworth, 1980; Fisher, 1931; Ponnikas et al., 2018; Wright et al., 2016), favouring the capture of beneficial alleles by recombination suppressors and accelerating sex chromosome divergence. More recently, it has been suggested that recombination arrest may instead be driven by the fixation of neutral suppressors (Ironside, 2010; Jeffries et al., 2021) or “lucky” ones that capture fewer than expected deleterious mutations (Jay et al., 2024; Lenormand and Roze, 2022). These theories have been partly motivated by the difficulty of directly observing sexual antagonism driving recombination evolution (Jay et al., 2024; Lenormand and Roze, 2022). Our results show that recombination suppression due to sexual antagonism evolves on a much faster timescale than that for neutral or lucky suppressors. Moreover, we find this is true even when sexual antagonism is sufficiently weak that it would be challenging to detect empirically. Beyond this, our analyses reveal a poorly appreciated pathway for sex chromosome evolution: recombination suppression may more often be driven by suppressors arising on the homogametic rather than the heterogametic chromosome. This bias should be especially pronounced in XY systems with high male reproductive variance, where selection on Y-linked suppressors shows diminished efficacy but selection on the X is largely unaffected. Testing these predictions empirically – especially in taxa where XY and ZW systems show different levels of recombination suppression – could help resolve the role of sexual antagonism in shaping sex chromosome evolution.

## Acknowledgements

We are grateful to the SNSF for funding (grants PCEFP3181243 and 310030220139) and to Sally Otto, Colin Olito, Denis Roze, Thomas Lenormand, Katie Peichel, and Andy Gardner for comments on a previous version and also to Colin Olito for clarifying the discrepancy between our results and those of Olito and Abbott (2025).

## Appendix A Evolution of polymorphism on the PAR

In this appendix, we detail gradual evolution at the trait locus on the PAR under the continuum of alleles model.

### A.1 Emergence of polymorphism

Under the model described in main text sections 2.1-2.2, gradual evolution at a locus under sex-specific selection has been studied with general fitness functions *w*_f_(*z*) and *w*_m_(*z*) in Flintham et al. (2025). They showed that selection first drives the population towards an equilibrium at *z*^∗^ *=* 2*x*^∗^. With Gaussian fitness (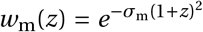and 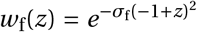), we can find this equilibrium by plugging *w*_m_(*z*) and *w*_f_(*z*) into eq. (A-11) in Flintham et al. (2025) and solving for *z*^∗^. This readily gives

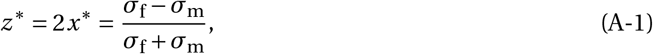

(it is also straightforward to see that this equilibrium is always convergence stable, i.e., is an attractor of evolutionary dynamics, by plugging in *w*_m_(*z*) and *w*_f_(*z*) into their eq. A-12).

Once the population expresses *z*^∗^, selection is then either stabilising, such that allelic values remain distributed around *x*^∗^ under mutation-selection balance; or disruptive whereby their distribution becomes bimodal in a process referred to as “evolutionary branching” leading to the maintenance of a polymorphism under balancing selection (Dercole and Rinaldi, 2008; Geritz et al., 1998; Rueffler et al., 2006). The condition for disruptive selection is found by plugging Gaussian *w*_m_(*z*) and *w*_f_(*z*) into eq. (E-27) of Flintham et al. (2025), which after some re-arrangements reads

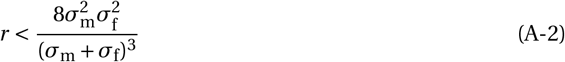

(see eq. B-25 in Flintham et al., 2025 for condition with general male and female fitness functions).

If condition (A-2) holds, the population will undergo evolutionary branching, leading to the emergence of two alleles (denoted a and A) that are held in a balanced polymorphism and that undergo further divergence (Geritz et al., 1998). We characterise this second stage of evolution, thus going beyond Flintham et al. (2025), in the following section.

### A.2 Divergence and maintenance polymorphism

The evolution of allelic values for a and A can be investigated through an invasion analysis that is based on the invasion fitness *W* (*x*_•_, *x*_a_, *x*_A_) of a rare mutant encoding an allelic value *x*_•_ in a population where resident A and a alleles – encoding *x*_a_ and *x*_A_, respectively – are at their equilibrium frequencies (Dercole and Rinaldi, 2008; Geritz et al., 1998; Otto and Day, 2011, chapter 12). The direction of the evolution of allelic effects a and A is determined by the selection gradient on each genetic value: *x*_A_ will evolve according to the sign of *s*_A_(*x*_A_, *x*_a_) *= ∂W* (*x*_•_, *x*_A_, *x*_a_)/*∂x*_•_|_*x*•*=x*A_ and *x*_a_ to the sign of *s*_a_(*x*_A_, *x*_a_) *= ∂W* (*x*_•_, *x*_A_, *x*_a_)/*∂x*_•_|_*x*•*=x*a_. Such evolution may therefore halt at an equilibrium allelic pair of values 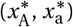 or “coalition” (Brown and Vincent, 1987) that satisfies

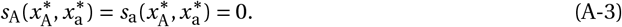

This coalition is convergence stable if the real part of the leading eigenvalue of the Jacobian matrix

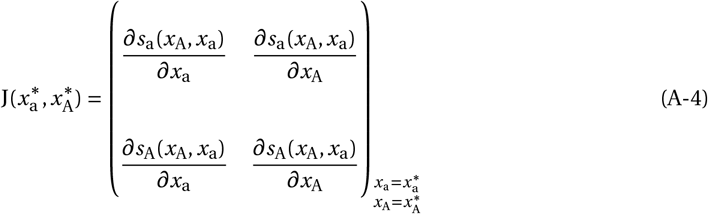

is negative, and is evolutionarily stable, i.e., there is no further branching once allelic values have converged to 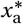 and 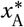, if

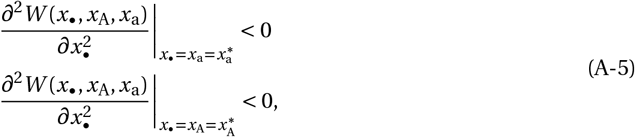

in which case the coalition 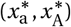 is an endpoint to gradual evolution.

#### A.2.1 Invasion fitness

We now characterise the invasion fitness *W* (*x*_•_, *x*_A_, *x*_a_) of a mutant encoding an allelic value *x*_•_ for our model. A mutant allele may initially appear in one of three chromosomal settings: it may arise (i) in an (necessarily X-bearing) egg cell, (ii) in an X-bearing sperm cell, and (iii) in a Y-bearing sperm cell. Its invasion fitness *W* (*x*_•_, *x*_a_, *x*_A_) is therefore given by the leading eigenvalue of the mean matrix

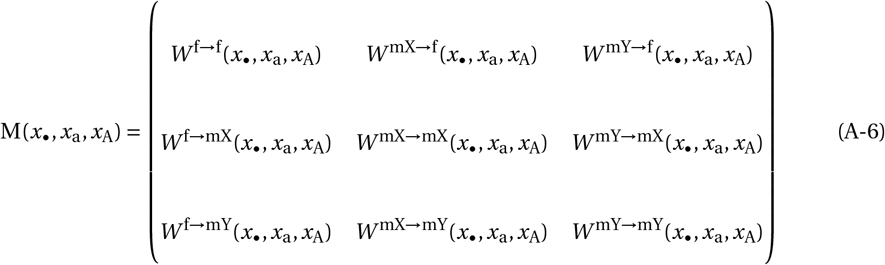

where *W* ^*u*→*v*^ (*x*_•_, *x*_a_, *x*_A_) is the expected number of recruited mutant gametes (i.e., that will go on to successfully form a surviving adult individual) of type *v* ∈ {f, mX, mY} at generation *t +*1 produced by a recruited mutant gamete of type *u* ∈ {f, mX, mY} at generation *t* (where superscript f denotes presence in eggs, mX in X-bearing sperm, mY in Y-bearing sperm).

It is straightforward to show that according to the life cycle of the population, the individual components of invasion fitness in eq. (A-6) are

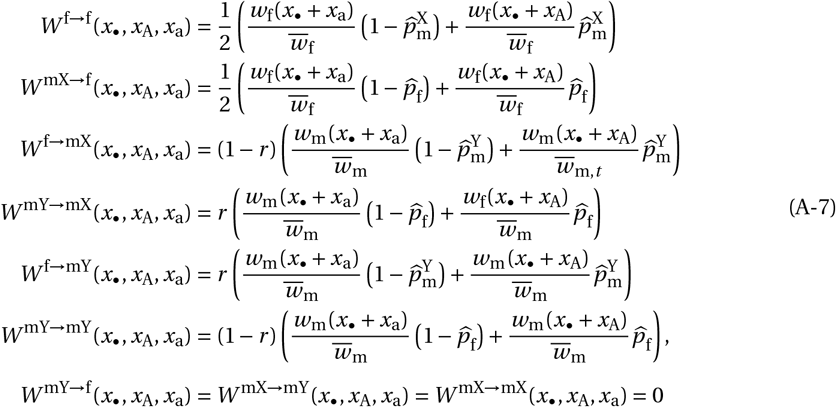

Where 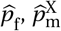, and 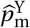 are the equilibrium frequencies of A in eggs, in X-bearing sperm, and Y-bearing sperm, respectively, and

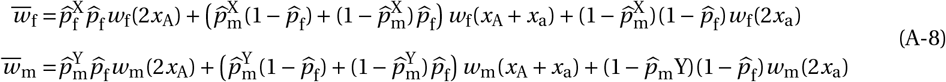

are mean female and male fitness when alleles A and a are at their equilibrium frequencies.

To find the equilibrium frequencies 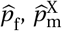, and 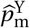, we write the frequency of allele A in each chromosome at generation at *t +* 1 as

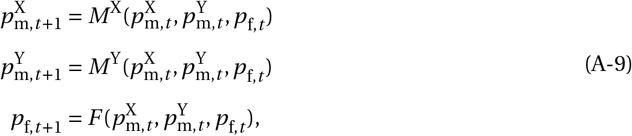

where

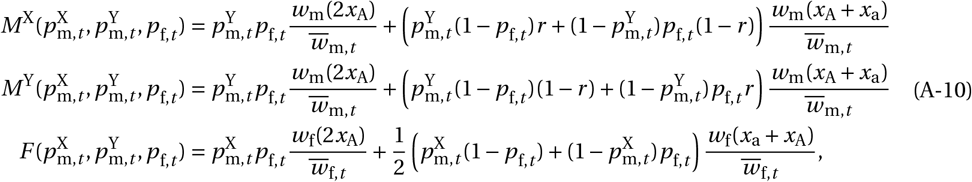

and

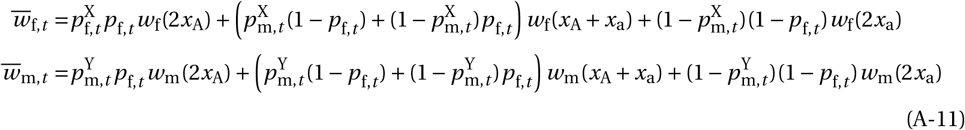

(Clark, 1988). Plugging eq. (A-10) into eq. (A-9), we can solve simultaneously for 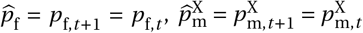, and 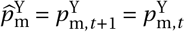.

#### A.2.2 Numerical approach

Owing to the complexity of the system eq. (A-9)-(A-11), we could not find analytical expressions for 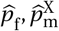 and 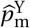. We therefore proceeded to find 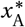 and 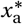 with a numerical analysis assuming Gaussian fitness functions for *w*_m_(*z*) and *w*_f_(*z*). To do so, we iterated the recursions

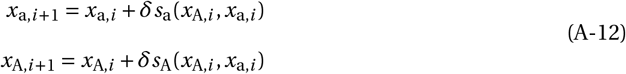

with *δ =* 0.1/((*σ*_m_ *+ σ*_f_)/2) (this scaling by *σ*_m_ and *σ*_f_ is made for computational expediency, so that changes in allelic values between iterations are not vanishingly small when selection is weak) until we reached a pair (*x*_A,*i*_, *x*_a,*i*_) that satisfied |*x*_A,*i+*1_ − *x*_A,*i*_ | < 10^−6^ and |*x*_a,*i+*1_ − *x*_a,*i*_ | < 10^−6^, which we took to be the equilibrium, i.e. 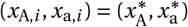. To calculate *s*_a_(*x*_A,*i*_, *x*_a,*i*_) and *s*_A_(*x*_A,*i*_, *x*_a,*i*_) at each iteration, we plugged *x*_a,*I*_ and *x*_A,*I*_ into eqs. (A-10) and eq. (A-11) to find the equilibrium frequencies 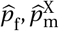 and 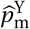, then plugged these allelic values and their associated equilibrium frequencies into eq. (A-7), then eq. (A-7) into eq. (A-6), from which we can calculate *s*_a_(*x*_a,*i*_, *x*_A,*i*_) and *s*_A_(*x*_a,*i*_, *x*_A,*i*_). Finally, we checked that the equilibrium 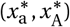 was convergence and evolutionarily stable by plugging these values into eqs. (A-4)-(A-5), which we found to be satisfied in all cases, indicating that 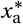 and 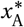 are stable endpoints to gradual evolution. We repeated this procedure with different parameter values (*σ*_m_, *σ*_f_, and *r*) to produce Fig 1C and SI Fig. 1. In each case, we initialised from a pair of weakly diverged allelic values (*x*_a,1_ *=* 0.01, *x*_A,1_ *=* −0.01).

### A.3 Balanced polymorphism in the PAR

In addition to the evolutionary stable values 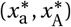, we also investigate arbitrary allelic values *x*_a_ and *x*_A_ that lead to a balanced polymorphism at the trait locus. For such a polymorphism for two alleles encoding *x*_a_ and *x*_A_ to exist requires that at least one eigenvalue of the Jacobian matrix

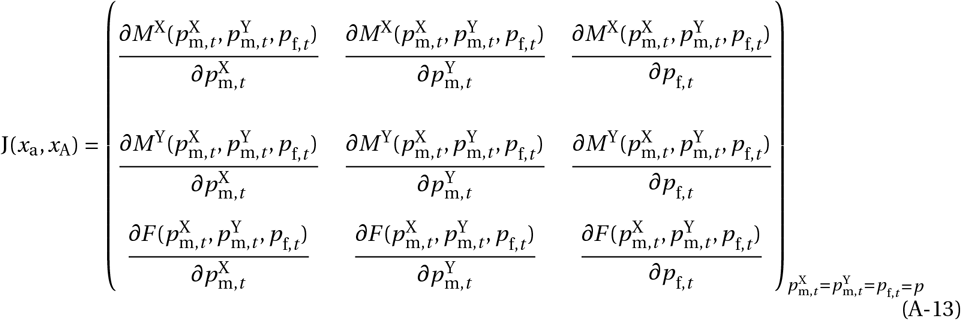

is greater than one in absolute value for both *p =* 0 and *p =* 1 (which indicates reciprocal invasion of alleles A and a), where the functions 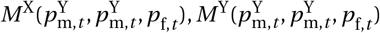 and 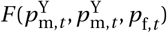 are detailed in eq. (A-10).

## B Simulating polygenic variation on the PAR

In this Appendix, we investigate whether male heterozygote advantage evolves in response to sexually antagonistic selection (as found under the continuum-of-alleles model, Appendix A) when *z* shows a polygenic basis on the PAR. To do this, we ran simulations where *z* was encoded by 10 di-allelic quantitative trait loci (with additive effects on the phenotypes) under the random-union-of-gametes model (Appendix C.3.1 for implementation of this lifecycle in a simulation). At each locus one segregating allele encodes an effect 0.25 and the other −0.25; these values were chosen for illustrative purposes, as they allow for a wide range of haplotype values, including the common evolutionarily stable values in the continuum-of-alleles-model 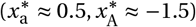. We assumed loci were equally spaced along the PAR with a uniform recombination rate *r* (i.e., *r* is the probability of recombination between adjacent trait loci, and between the SDR and the closest terminal trait locus) and we initialised simulations with equal frequencies of each allele at each locus. We tracked the evolution of the genetic values encoded by X and Y chromosomes over 15000 generations of sexually antagonistic selection (under Gaussian fecundity functions with *σ =* 0.1 for illustration). Results are shown in Appendix Fig. 1.

**Appendix Figure 1:**
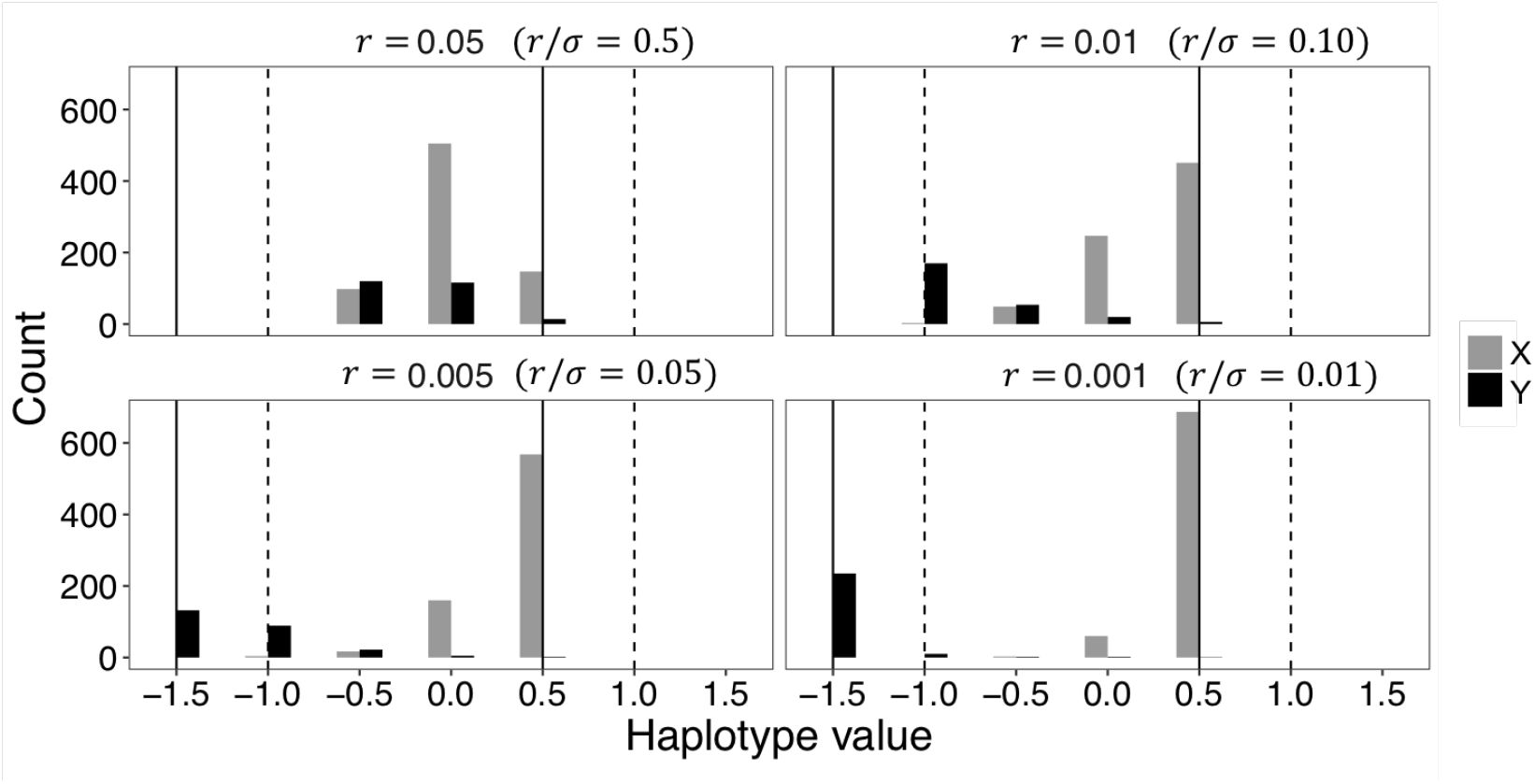
Evolved haplotypes when *z* is polygenic. Panels show distributions of genetic effects encoded by a sample of 1000 chromosomes (grey and black columns for X and Y chromosomes, respectively) segregating in the population after 15000 generations of a simulation. Different panels show results for different strengths of recombination amongst loci, *r*. Vertical dotted lines indicate the location of the male and female optima (at −1 and 1, respectively) and vertical solid lines indicate the evolutionary stable allelic values (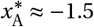 and 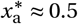, see Fig. 1C).

With strong recombination (e.g., *r* /*σ =* 0.5), genetic values for the X and Y evolve to be distributed symmetrically around 0 (similar to autosomal chromosome pairs, Flintham et al., 2025), however, where recombination is weaker (e.g., *r* /*σ =* 0.1 or *r* /*σ =* 0.05), X-chromosome haplotypes develop that tend to encode an allelic value of 0.5 or smaller, while Y-chromosome haplotypes encode a much larger absolute effect that overshoots the male optimum in homozygous state. Indeed when linkage is tight, e.g., *r* /*σ =* 0.01, haplotype values are distributed around the evolutionary stable values (i.e., X-haplotype encoding 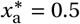 and Y-linked haplotype encoding 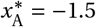). Consequently, when *z* shows a polygenic basis that is in linkage with the SDR (i.e., in the PAR), we expect that females show highest fitness when homozygous for the most common X-linked haplotype, and so are homozygotes across trait loci. Meanwhile, the highest fitness males, that is, those carrying one copy of the most common Y- and X- haplotypes, are heterozygous across loci, leading to heterozygote advantage.

## C. Evolution of a Y-linked suppressor

### C.1 Invasion fitness

Here, we characterise the invasion fitness of a Y-linked recombination suppressor appearing as a single copy in a recruited Y-bearing sperm (i.e., that successfully goes on to form an adult male). This mutant appears arises in a population that recombines at a resident rate *r* and that is at a polymorphic equilibrium at the trait locus – i.e., where alleles A and a are at their equilibrium frequencies given by 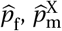 and 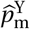. Such a mutant arises in a Y-bearing sperm carrying allele A with probability 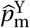 and one carrying allele a with probability 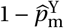. Let us denote with 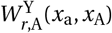 and 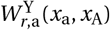 the invasion fitness for each of these two cases respectively. Since we assume the suppressor fully arrests recombination with the trait locus, these are given by

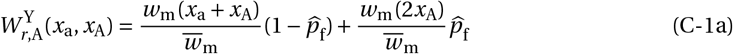

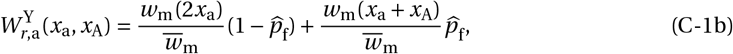

where recall 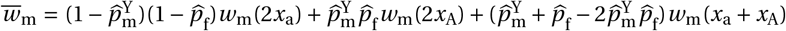 is mean male fitness with allele frequencies (see also Olito and Abbott, 2025 eq. 15 and Appendix C for an alternative derivation of an expression for a di-allelic sexually antagonistic locus with fixed effects, and Otto, 2014 for expressions for a small effect recombination modifier).

For a suppressor to invade given it arises with allele A or with allele a, we must have 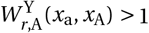 and 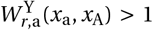, respectively. We first argue that 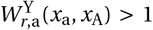 never holds. We do so by noting first that once alleles A and a have diverged (i.e., *x*_A_ < *x*_a_), additive gene action entails that male heterozygotes show higher fitness than those homozygous for allele a (i.e., *w*_m_(*x*_a_ *+ x*_A_) > *w*_m_(2*x*_a_)). Allele A can then either : (I) be strictly male-beneficial: *w*_m_(2*x*_a_) < *w*_m_(*x*_a_ *+ x*_A_) < *w*_m_(2*x*_A_), or (II) lead males to show heterozygote advantage: *w*_m_(*x*_a_ *+ x*_A_) > *w*_m_(2*x*_a_) and *w*_m_(*x*_a_ *+ x*_A_) > *w*_m_(2*x*_A_). In scenario (I), with some rearrangements of eq. (C-1b), it is straightforward to show that 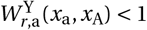 for all 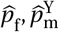 and all *x*_a_ and *x*_A_ (with *x*_A_ < *x*_a_), so that a suppressor capturing allele a cannot invade. In scenario (II), rearranging eq. (C-1b) shows a suppressor can invade if the following condition holds

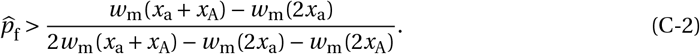

This requires the equilibrium frequency of allele A be large relative to the fitness advantage of male heterozygotes. We evaluated this condition for a large range of allelic values *x*_a_ and *x*_A_ satisfying *w*_m_(*x*_a_ *+ x*_A_) > *w*_m_(2*x*_A_) and that would result in a balanced polymorphism at the trait-locus (Appendix A.3; including the equilibrium 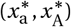), strengths of selection (*σ*_m_ and *σ*_f_), and recombination rates *r* numerically (using eqs. A-9-A-10 to find the equilibrium frequencies for alleles A and a, and inserting these values into eqs. C-5b and C-2). Among 1000 parameter sets tested in *σ*_m_ ∈ [0.0001, 1], *σ*_f_ ∈ [0.0001, 1], *r* ∈ [0.0025, 0.5], *x*_a_ ∈ [0.01, 0.5], *x*_A_ ∈ [−0.01, −1.6] we found none such that condition (C-2) is fulfilled.

By contrast, using the same arguments, we find that 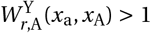 always holds: When A is male-beneficial (scenario I above), it is straightforward to show that 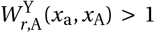 is satisfied for all 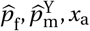, and *x*_*A*_; main text condition (C-5a). Whereas, if males show heterozygote advantage (scenario II above), then 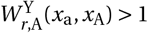 when

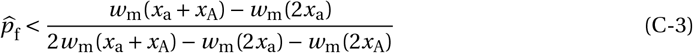

(given as main text condition C-5b – eq. C-4 is the mirror image of eq. C-4). Performing the same numerical analysis as for condition eq. (C-2), we found that eq. (C-3) was fulfilled for all 1000 parameter sets we tested in *σ*_m_ ∈ [0.0001, 1], *σ*_f_ ∈ [0.0001, 1], *r* ∈ [0.0025, 0.5], *x*_a_ ∈ [0.01, 0.5], *x*_A_ ∈ [−0.01, −1.6].

Having established that the capture of allele A is both a necessary and sufficient condition for invasion of a Y-linked suppressor, to simplify notation we drop the dependency of 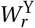 on allele type and use it to refer to the invasion fitness of a recombination suppressor conditional on it capturing allele A, i.e., 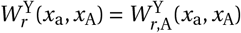 (this notation is also used in the main text).

We next investigated how the strength of positive selection on a suppressor capturing allele A, measured by the magnitude of invasion fitness (i.e., how much greater than unity 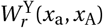 is), depends on allelic values (*x*_a_ and *x*_A_), the strength of Gaussian sex-specific selection (*σ*_m_ and *σ*_f_), and the recombination rate (*r*). As before, we used eqs. (A-9)-(A-10) to solve for 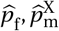 and 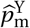 for a wide range of parameter sets that precipitate a balanced polymorphism at the trait locus (Appendix A.3), and plugged these frequencies and their associated parameters into eq. (C-1a) in order to produce Fig. 2. These results are summarised and interpreted in the following section (Appendix C.2).

### C.2 Sexual antagonism favours recombination suppression, especially with evolutionarily stable polymorphism

In Appendix C.1 we showed that a Y-linked suppressor of recombination can invade due to selection only if it captures the male-beneficial allele A (where *x*_A_ < *x*^∗^ < *x*_a_), in which case its invasion fitness reads as

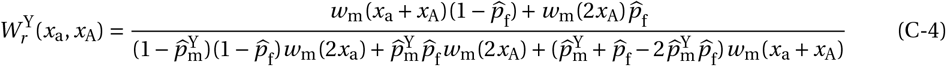

(eq. C-1b written out in full – although the resident recombination rate *r* does not appear explicitly in eq. (C-4), *r* does influence 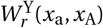 indirectly because the equilibrium frequencies of A depend on it; i.e., 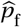 and 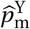 depend on *r, x*_A_ and *x*_A_). A recombination suppressor can be favoured by selection 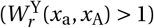 via two routes: either

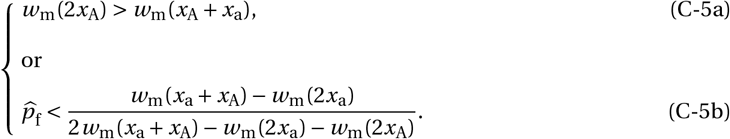

The first route (eq. C-5a) is for the homozygote AA to encode the highest fitness in males (i.e. *w*_m_(2*x*_A_) > *w*_m_(*x*_A_ *+ x*_a_)), whereas the second route (eq. C-5b) applies if instead the heterozygote Aa encodes the highest fitness (i.e. *w*_m_(*x*_A_ *+ x*_a_) > *w*_m_(2*x*_A_) such that males exhibit heterozygote advantage). The increased certainty in the patrilineal transmission of allele A brought by recombination suppression is advantageous if either: (a) this allele A is systematically male-beneficial such that AA homozygotes enjoy the greatest male fitness (as in condition C-5a); or (b) males show heterozygote advantage and the frequency 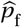 of allele A amongst eggs is sufficiently low such that a male always passing on A through his Y-chromosome produces male offspring that are mostly Aa heterozygotes (hence condition C-5b being more stringent when 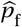 is large).

Computing 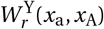 across a range of parameter values shows that provided a and A are maintained by balancing selection in the resident population, selection always favours a full recombination suppressor, i.e., 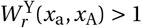 (as known to be the case for a small effect recombination modifier; Otto, 2014, see her condition for selection to favour suppression with *R*_*m*_ *=* 0). For weakly diverged alleles (*x*_a_ and *x*_A_ similar), this is simply because condition (C-5a) is always satisfied when a and A segregate. For alleles that generate male heterozygote advantage, this tells us that under conditions leading to balancing selection on the PAR, recombination and sexually antagonistic selection always drive sufficiently low frequencies 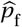 of allele A in eggs to satisfy condition (C-5b).

Our results also indicate that the strength of selection on suppression is sensitive to the strength of recombination in the resident population, with 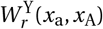 being substantially greater when *r* is large (compare Figs. 2A-C; see also Olito and Abbott, 2025). This is because when *r* is small, the frequency 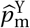 of A in Y-bearing sperm is high anyway, which diminishes the benefit to a mutant male that ceases recombination.

Selection for recombination suppression is especially strong when alleles have evolved to their evolutionary equilibrium (i.e. 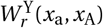 is especially large when 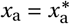 and 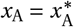, Figs. 2A-C, white stars). This is due to two complementary effects. First, the equilibrium values 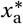 and 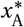 entail a large variance in male fitness under random mating, owing to their substantial divergence (in general, selection for recombination suppression increases with the divergence between *x*_a_ and *x*_A_, and so is greatest under the conditions leading to male heterozygote advantage, Fig. 2). Thus, with equilibrium allelic values, Aa males experience an especially large relative fitness advantage over homozygote genotypes, which in turn leads to a strong fitness benefit to be conferred by a recombination suppressor that ensures a male’s sons inherit A on their Y-chromosome. Second, the equilibrium value 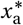 generates the strongest possible positive selection on allele a through females (due to strong homozygote aa advantage), and so leads to the lowest equilibrium frequencies of allele A on the X-chromosome (i.e., low values of 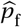 and 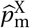). This means resident males on average carry an excess of allele a via their maternal X, and so depart more frequently from the male optimum, reinforcing the competitive advantage of males that carry allele A on their Y-chromosome.

### C.3 Probability of invasion and substitution times under random mating

#### C.3.1 Probability of invasion

As a next step in our analysis, we calculate the probability Ψ^Y^ that a Y-linked suppressor that has arisen as a single copy successfully invades, i.e. evades stochastic loss. From Appendix C.1 and standard branching process theory, we know that a Y-linked suppressor is only under positive selection when it captures allele A; otherwise the suppressor experiences negative selection and so goes extinct with probability 1 when it arises as a single copy. The (unconditional) invasion probability of a suppressor can therefore be expressed as 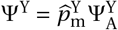. Here, 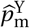 is the frequency of allele A in Y-bearing sperm, and so gives the probability a Y-linked suppressor captures allele A, and 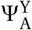 is the (conditional) probability a suppressor invades given it captured allele A.

This conditional invasion probability can be calculated using a branching process argument (Haldane, 1927; Otto and Whitlock, 2013) by solving

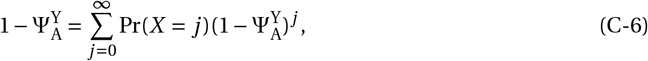

where *X* is a random variable denoting the number of copies produced by a rare suppressor that has captured allele A over one generation (such that Pr(*X = j*) for *j* ∈ {0, 1,…} gives the probability mass function of suppressor fitness). Equivalently *X* is a random variable for the number of surviving male offspring produced by a male carrying the suppressor mutation.

Under the lifecycle given in section 2.1 (i.e. the random-union-of-gametes), *X* follows a Poisson distribution with mean given by 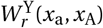 (shown in eq. C-1a), i.e. *X* ∼ Pois 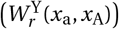 so that

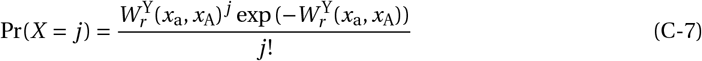

(we consider alternative mating system, leading to a different form for Pr(*X = j*) in Appendix C.3.3). Substituting eq. (C-7) into eq. (C-6), we obtain

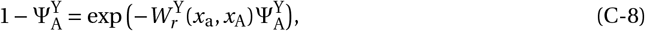

which cannot be solved in general.

To generate our main text results, we take a numerical approach and calculate Ψ^Y^ across a wide range of allelic values *x*_a_ and *x*_A_ (including the equilibrium values 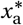 and 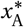), strengths of Gaussian sex-specific selection *σ*_m_ and *σ*_f_, and resident recombination rate *r*. This involved (1) calculating the equilibrium allele frequencies 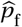 and 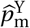 and invasion fitness 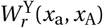 (numerically as per Appendices A.2.1 and C.1) for a given allelic pair of values (*x*_a_, *x*_A_) and parameter set (*σ*_m_, *σ*_f_ and *r*); (2) plugging 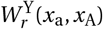 into eq. (C-8) and solving numerically for the conditional invasion probability 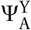; (3) multiplying 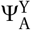 by 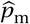 to give the unconditional probability Ψ^Y^. Our results for Ψ^Y^ are shown in Fig. 3A and SI Fig. 4 and interpreted in main text section 3.2.

Our analysis here can be connected to that of Olito and Abbott (2025), who also calculate the unconditional invasion probability 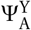 for a Y-linked recombination suppressor that captures a sexually antagonistic locus with fixed additive fitness effects (specifically where *w*_f_(2*x*_a_) *=* 1, *w*_f_(*x*_a_ *+x*_A_) *=* 1−*s*_f_/2, *w*_f_(2*x*_A_) *=* 1 − *s*_f_, *w*_m_(2*x*_a_) *=* 1 − *s*_m_, *w*_m_(*x*_a_ *+ x*_A_) *=* 1 − *s*_m_/2, *w*_m_(2*x*_A_) *=* 1 and 0 ≤ *s*_m_ < 1 and 0 ≤ *s*_f_ < 1). Olito and Abbott (2025) use eq. (C-1a) to calculate 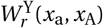 for a Y-suppressor capturing the strictly male-beneficial allele A and then use the approximation 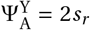 where 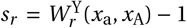. This works well when selection on a modifier is weak (*s*_*r*_ is small; e.g., under the branching process, writing 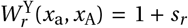, where *s*_*r*_ ≪ 1, gives 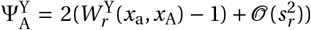, but strongly overestimates invasion probabilities when selection is strong (Otto and Whitlock, 2013), and so is more limited for studying recombination suppression when polymorphism evolves at PAR loci far from the SDR-boundary (*r* and *σ* large), as we do here.

#### C.3.2 Fixation and substitution

Our results thus far have concerned the dynamics of a suppressor as it invades into a population from a single copy. However, the tempo of recombination evolution depends not just on how frequently such suppressors invade, but also on whether they go on to fix in the population, the conditions for which may diverge from those for invasion. In particular, a large effect mutation such as a full recombination suppressor (i.e., that completely silences recombination) that invades into a population may be prevented from fixing if it experiences negative frequency-dependent selection – in which case the suppressor is held at an intermediate frequency as a balanced polymorphism.

To establish whether a Y-linked suppressor that has successfully invaded into a population will go on to fix or not, we ran individual-based simulations of a Wright-Fisher process using SLiM 4,0 (Haller and Messer, 2023) that track the frequency of a suppressor through time. Following the lifecycle in section 2.1, we simulate a diploid population of *N =* 10^4^ adults with non-overlapping generations, where each individual carries a single chromosome with three loci: one XY sex-determining locus, one modifier locus that is in full linkage with the sex-determining locus, and one trait locus recombining with the sex-determining locus at rate *r*). At each generation, 5000 male and 5000 female individuals are created through multinomial sampling of adult males and females from the previous generation, with a sample weighting given by their fecundity (*w*_m_(*z*) and *w*_f_(*z*), which as we assume are Gaussian, where *z* is given by the sum of the values encoded by the alleles they carry at the trait locus). We assume the trait locus is polymorphic for alleles A and a and we initialise simulations with alleles at their equilibrium frequencies in all gene pools (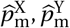and 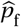; to ensure one allele is not lost by chance during the simulation, we allow mutations between the two alleles at a per-locus rate 5 *×* 10^−5^).

At the beginning of each simulation, we introduce a suppressor mutant into the population, doing so in one of two different ways: (1) In one set of simulations, we introduce a single copy into the population (i.e., with initial frequency 1/(*N* /2)), which is placed in the Y-chromosome of a randomly sampled male. This setup therefore follows our previous analyses in considering the fate of a novel suppressor (providing a comparison for our invasion results). (2) In a second set of simulations, we introduced the mutant into *q N* /2 (rounded to the nearest integer) randomly sampled A-bearing adult male Y-chromosomes, where 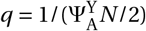 is the frequency at which a mutant suppressor that originally arose a single copy in a chromosome containing A can be considered to have invaded (i.e., to have avoided stochastic loss when rare in a large population, Charlesworth, 2020; Maynard Smith, 1976). In other words, here we are following the trajectory of a suppressor, conditional on it successfully establishing in the population. In both cases, we ran simulations until the suppressor either fixed or went extinct in the population, doing so for the same range of parameter/allelic value combinations considered in section C.3.1, with 10^4^ replicates for each parameter set.

For all parameter sets considered we found: (1) In simulations where a suppressor was introduced as a single copy, the proportion of replicates in which the mutant fixed closely matched the unconditional invasion probability, Ψ^Y^, calculated in Appendix C.3.1 (some example values are plotted in Fig 3A, and the frequency trajectories of some example simulations are shown in Appendix Fig. 2A). (2) in simulations where the mutant was introduced at frequency *q*, we found that it always quickly swept to fixation. Together, these observations strongly suggest that successful invasion by a Y-suppressor is sufficient for its eventual fixation (i.e., selection on recombination suppressors is not frequency dependent even when selection is strong; this is consistent with the results of Olito and Abbott, 2025 Appendix C, and extends the weak selection results of Otto, 2014 who found that the invasion conditions for small effect recombination promoters and suppressors are mutually exclusive). Consequently, we consider that Ψ^Y^ captures both the probability of invasion and fixation for a Y-linked suppressor.

**Appendix Figure 2:**
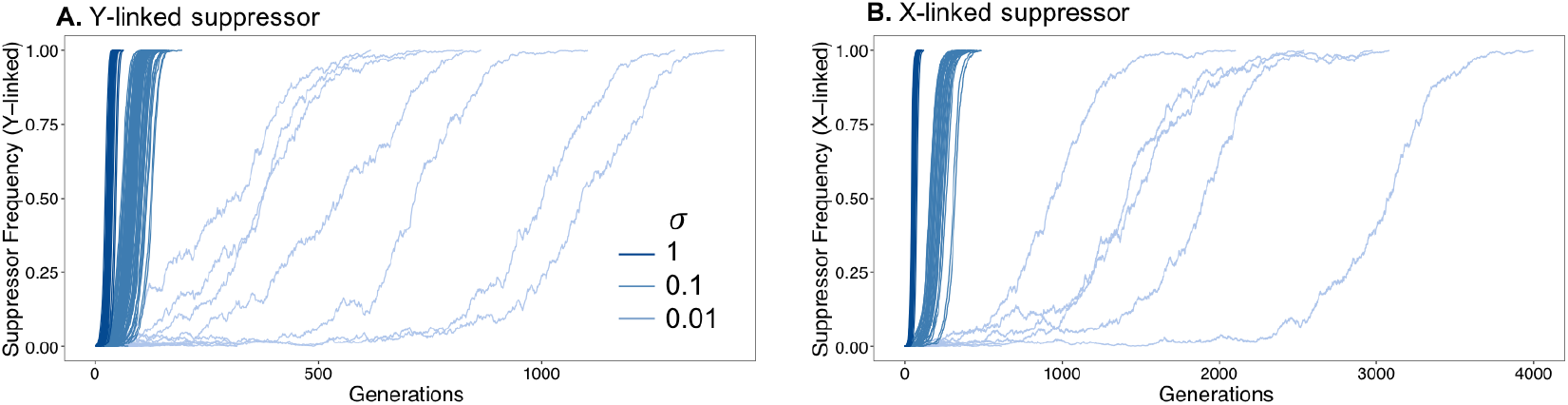
Frequency of Y- and X-linked suppressors through time from example simulations. Plots show suppressor frequency through time in example simulations where a suppressor was introduced as a single copy (Appendix C.3.2 and D.2.2 for details). Panels **A**. and **B**. respectively show frequency trajectories for simulations where a Y-linked and X-linked suppressor was introduced and different colour curves show replicates with different strengths of sexually antagonistic selection (*σ = σ*_f_ *= σ*_m_, other parameters: *x*_a_ *=* 0.5, *x*_A_ *=* −0.5, *r =* 0.5). A sample of 1000 replicate simulations is shown for each treatment, with two mutually exclusive types of trajectories being observed for both Y- and X-linked suppressors: (1) suppressor frequency remains low after its introduction, and the mutant ends up being lost quickly, (2) suppressors rapidly sweep to fixation at a rate sensitive to *σ* (notice curve show the characteristic sigmoid-shape consistent with a sweep occurring). This pattern supports the notion that suppressors that successfully invade into a population systematically go on to fix.

With this in mind, we can use our invasion results to calculate the expected waiting time for such a suppressor to substitute (to appear in a population and then go on to fix). To do this, we assume suppressor mutants arise in a population at a low constant rate *λ*_*µ*_ *=* (*N* /2)*µ* ≪ 1 per generation (where *µ* is the probability a novel suppressor mutant appears in a given Y-bearing gamete), such that substitution is a Poisson process with rate *λ*_sub_ *= λ*_*µ*_Ψ^Y^. From this it follows that the random variable *T* giving the waiting time (number of generations) for the substitution to occur is exponentially distributed *T* ∼ Exponential(*λ*_sub_), and therefore with mean

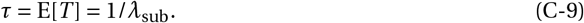

To generate our main text Fig. 3B we plugged the values for Ψ^Y^ calculated numerically into eq. (C-9), which we compared against the neutral substitution time: 1/*µ*. In our figures we used a large population size 10^4^ and a gametic mutation rate of 10^−5^ (as in Lenormand and Roze, 2022), although equivalent results are produced with weaker mutation rates. Recent estimates (Berdan et al., 2021, box 1) place genome-wide gametic mutation rates for recombination suppressors such as inversions at ≈ 10^−3^ (and chromosomal fusions at ≈ 10^−4^), thus our parameters are consistent with 1% of these mutations occurring on the Y-chromosome and capturing the SDR.

#### C.3.3 Probability of invasion under polygyny

Here, we consider the invasion probability Ψ^Y^ under a model of lottery polygyny (Nunney, 1993). To do so, we can still use eq. (C-6) but *X* should now be non-Poisson. We determine the probability mass function Pr(*X = j*) under a lifecycle similar to the one analysed by Lee et al. (2008). This life-cycle is as follows : (1) *N* /2 adult males and *N* /2 adult females enter the mating pool (where *N* is large). Females mate singly but males can mate multiply. For each female, a male is sampled with replacement which mates with her and sires all her offspring. The probability that a male *i* ∈ {1,…, *N* /2} is sampled is weighted by his competitiveness which we denote as *c*_*i*_ (e.g., *c*_*i*_ represents a male’s biological condition, or level of expression of a sexually selected ornament relative to other males in the population). (2) After mating, females produce a large number of juveniles (in this model, male mating success and female fecundity are independent of genotype). (3) Adults die and juveniles undergo sex-specific density-dependent competition for access to *N* /2 male and *N* /2 female adult positions in the mating pool of the next generation. This competition is mediated by the evolving trait *z*; i.e. the relative probability of surviving to the mating pool of a juvenile male expressing trait *z* is *w*_m_(*z*), while that of a female is *w*_f_(*z*). This ensures that the expected number of surviving offspring of males and females expressing a trait value *z* is identical to that in the random-union-of-gametes model (section 2.1) and therefore leads to equivalent sexually antagonistic selection in both models, allowing us to focus on the effects of mating system on reproductive variance.

For this model of polygyny, we can obtain the probability mass function Pr(*X = j*) for the number *X* of males produced by a mutant male using conditional probabilities :

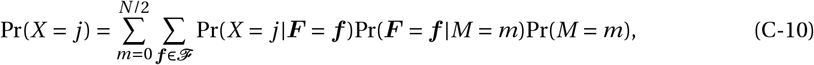

where *M* is the random variable for the number of females mated by a mutant male, and ***F*** *=* (*F*_aa_, *F*_Aa_, *F*_AA_), is a vector containing the random variables for the numbers of females of each genotype the mutant male mates (e.g., *F*_aa_ is a random variable for the number of aa females he mates).

By definition, *M* has support {0, 1,…, *N* /2}, and if male competitiveness follows a Beta distribution with parameters *α* and *β = α*(*N* /2−1) (such that *C*_*i*_ ∼ B(*α, α*(*N* /2−1)) where *C*_*i*_ is the random variable for competitiveness for each male *i*), then

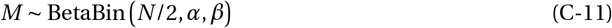

where BetaBin refers to the Beta-binomial distribution (Lee et al., 2008, eq. 1 with their *n*_*i*_ large). The random-variable *M* has expectation E[*M*] *=* 1 and variance Var[*M*] *=* (*N* −2)(1*+α*)/(2*+Nα*) → 1*+*1/*α* as *N* → ∞. The parameter *α* therefore scales the variance in male mating success (through its effect on the distribution of male competitiveness), which is high when *α* is small and low when *α* is large.

Because mating is random with respect to genotype, the number of females of each genotype (AA, Aa, and aa) that the mutant male mates with, given he mates a total *m* females follows a Multinomial law with parameters

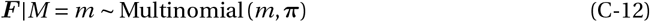

where 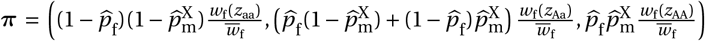 gives the frequencies of each genotype amongst adult females. We have denoted the support of this random variable as 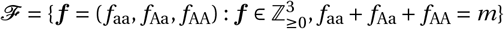.

Finally, given it mates with a group of ***f*** *=* (*f*_aa_, *f*_Aa_, *f*_AA_) females, the number of surviving male offspring a mutant male sires (the random variable *X*) is the sum of all the surviving sons of these females. According to our lifecycle (whereby females each produce a large number of juveniles that each compete independently and sex-specifically to obtain a breeding spot), the numbers of surviving male offspring of each genotype produced by a given female follow poisson distributions with means equal to the relative male fitness of each genotype weighted by the proportion of the focal female’s juvenile offspring that will carry each genotype. For example, a female with genotype Aa mated to a male carrying A on his Y-chromosome will produce AA and Aa juvenile sons in equal proportions, and so will leave Poisson distributed numbers of surviving male AA and Aa offspring with means 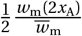 and 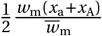, respectively. Putting all this together, we have

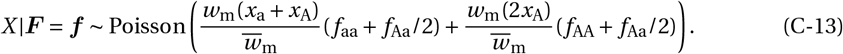

Using eqs. (C-11)-(C-13) for eq. (C-10) then gives us the unconditional probability mass function Pr(*X = j*).

To generate main text Fig. 3C we plugged eq. (C-10) into eq. (C-6) and solved numerically for 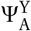, doing this for different values of *α* that lead to different mating variances. For computational expediency we performed the calculation with small population sizes; *N =* 20 when *α* ≥ 1 and *N =* 100 otherwise, but which were nonetheless sufficient to capture full probability distribution of *M*, i.e., Pr(*M* ≤ 10) ≈ 1 when *α* ≥ 1 and Pr(*M* ≤ 50) ≈ 1 for all other values of *α* considered, so that this procedure was not problematic given our broader assumption that *N* is large. We calculated substitution times in the same way as for the random-union-of-gametes model (described in Appendix C.3.2) to produce Fig. 3D.

In addition, we ran individual-based simulations for the fixation of a recombination suppressor under lottery polyny to compare with our numerical results for Ψ^Y^. To apply the polygyny lifecycle, we modelled a population of fixed size *N* with *N* /2 male and female adults with the following stages: Each generation, adult females independently sample a mate (i.e., males are sampled with replacement) to sire all her offspring. A males’ sampling probability was weighted by his competitiveness, drawn from a Beta distribution. Females then each produce 20 juveniles. Adults die and *N* /2 juvenile males and *N* /2 juvenile females are sampled (without replacement) to become the adults of the next generation, while all other juveniles die. A male juvenile’s sampling probability was weighted by *w*_m_(*z*) and a female juvenile’s by *w*_f_(*z*). Following the same simulations procedure as in Appendix C.3.2, we computed proportion of simulations in which a novel suppressor fixed when introduced as a single copy, and found a good match with the unconditional invasion probability, Ψ^Y^, calculated using eq. (C-10) (see dots in Fig. 3C).

## D Evolution of an X-linked suppressor

### D.1 Invasion fitness

In this Appendix we consider the fate of a new modifier on the X-chromosome that arises in the SDR-homologue and suppresses recombination with the trait locus in male or female heterozygotes for the modifier (as per a chromosomal inversion). Because a new X-linked suppressor may arise in either a male or a female chromosome, we compute its invasion fitness, 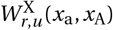, conditional on capturing allele *u* ∈ {a, A}, as the leading eigenvalue of the mean matrix

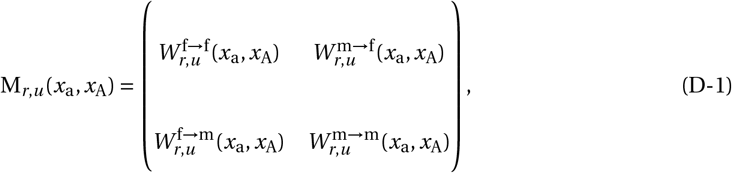

where 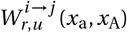 gives the number of recruited gametes of sex *j* ∈ {f, m} (i.e., number of eggs and number of sperm, respectively) containing allele *u* ∈ {a, A} produced over the course of a generation by a recruited gamete of sex *i* ∈ {f, m} and carrying allele *u* ∈ {a, A}. These fitness components are:

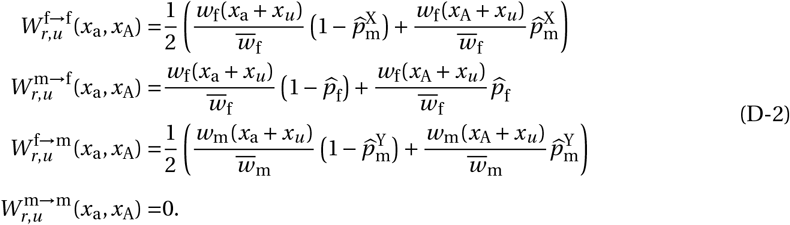

where 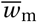 and 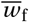 are given by eq. (A-8).

The quantities 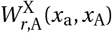 and 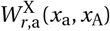 end up being cumbersome, and so are given in a Supplementary Mathematica Notebook. They also do not lead to readily interpretable invasion conditions (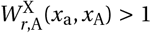 and 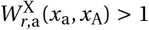) for an X-linked modifier, so we again proceed with a numerical analysis. Using Gaussian functions for *w*_f_(*z*) and *w*_m_(*z*) and following an identical procedure as in Appendix C.1, we find that across all parameter (*σ*_m_, *σ*_f_, *r*) and allelic value (*x*_a_, *x*_A_) combinations tested, an X-linked suppressor invades only when it captures the female-beneficial allele a, and is otherwise disfavoured by selection (i.e., 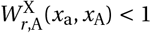 and 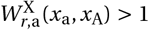 for all *x*_a_, *x*_A_, *σ*_m_, *σ*_f_ and *r*). We therefore drop the dependency of invasion fitness (and its components) on allele *u*, and use the notation 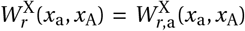 in the main text (e.g., in section 3.3) and for the rest of the Appendix. We present plots of 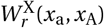 for different allelic values and parameters in SI Fig. 5 (calculated with the same procedure as Appendix C.1), and the strength of selection for a Y-suppressor relative to an X-suppressor 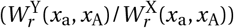 in SI Fig. 6A.

### D.2 Invasion and fixation probabilities for X-linked suppressors based on multitype branching process

#### D.2.1 Probability of invasion

As an X-linked suppressor can only be favoured by selection when it captures allele a, its unconditional probability of invasion, Ψ^X^, depends on its probabilities of invasion conditional on the mutant arising on a male, 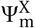, and female, 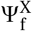, X-chromosome carrying allele a, weighted by the equilibrium frequencies of this allele; 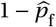 and 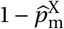. Specifically, taking into account the difference in male and female X copy numbers (and so likelihood of a given X-linked mutation first appearing in a male or female), we have

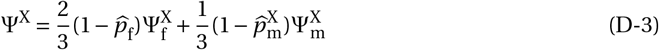

(Charlesworth et al., 1987).

The growth of the mutant suppressor arisen as a single copy follows a multitype branching process (Harris et al., 1963). As such, if a rare mutant can invade into a population (i.e., if 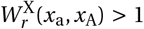), the probability this mutant ultimately invades conditional on it arising in a gamete produced by sex *i* ∈ {f, m} is 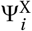. If mutant fitness distributions are Poisson for all genetic backgrounds, that is, if 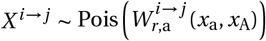 for all *i* ∈ {f, m} and *j* ∈ {f, m}, where *X* ^*i*→*j*^ is the random variable for the number of mutants in a gamete of sex *j* produced by a mutant in gamete of sex *i*, then 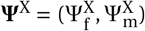 is the unique solution to the set of simultaneous equations

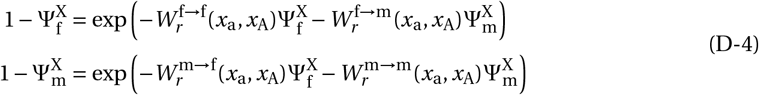

(where recall 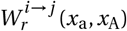 refers to 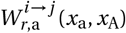 in eq. D-2). Taking the log of both sides of eq. (D-4) and rearranging, we arrive at the system

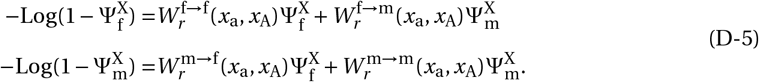

After plugging in eq. (D-2), eq. (D-5) can be solved numerically for 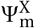 and 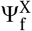 using the same approach as we used to find Ψ^Y^ in Appendix C.3.1. Finally, plugging these quantities into eq. (D-3) with the corresponding equilibrium frequencies 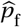 and 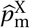 allowed and comparing with the equivalent invasion probabilities for Y-linked suppressors (to calculate Ψ^Y^/Ψ^X^ and *ρ =* Ψ^Y^/(Ψ^Y^ *+* Ψ^X^)), we could produce Fig. 4A and SI Fig. 6B.

#### D.2.2 Fixation and substitution

In order to confirm that successfully invading X-linked suppressors also go on to fix, we ran individual-based simulations following the same procedure as in Appendix C.3.2. Depending on the scenario being considered, we either introduced a suppressor (1) as a single copy into the population, by placing it in a randomly sampled X-chromosome (which could be present in an adult male or female; example frequency trajectories are shown in Appendix Fig. 2B), or (2) at a frequency *q =* 1/(3*N* /4Ψ^X^) by adding it to 3*Nq*/4 (again rounded to the nearest integer) randomly sampled abearing X-chromosomes. Across the same parameter sets as used in Appendix C.3.2, we found that when introduced at frequency *q*, X-linked suppressors always went on to fix in the population. Furthermore, for simulations where the suppressor was introduced as a single copy, we saw a good match between the proportion of replicates in which this mutant fixed and our numerical calculation of Ψ^X^ in Appendix D.2.1 (see dots in Appendix Fig 3). This is consistent with invasion implying fixation for X-linked suppressors and we therefore interpret Ψ^X^ as both the probability of invasion and fixation for our parameter sets.

**Appendix Figure 3:**
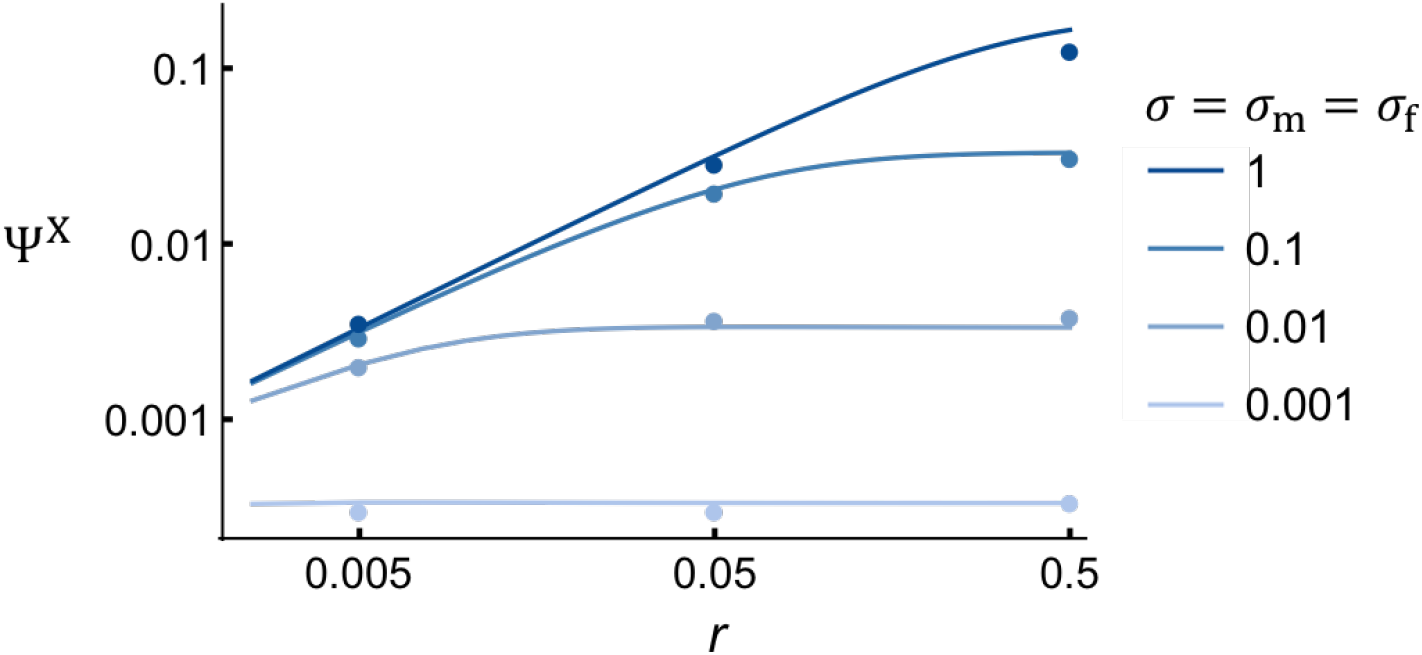
Invasion of X-linked suppressors in a large population. Figure shows the unconditional invasion probability of a newly arisen X-linked suppressor in a large population, Ψ^X^, under the random-union-of-gametes model (Appendix D.2.1 for calculation) according to the resident recombination rate (*r*). Allelic values used: *x*_a_ *=* 0.5 and *x*_A_ *=* −0.5. Dots show results from individual-based simulations with (which correspond to proportion of replicate simulations in which a X-suppressor fixed, Appendix D.2.2). Different colour curves and dots correspond to results for different strengths of sexually antagonistic selection at the trait locus (*σ = σ*_m_ *= σ*_f_).

In contrast to these results, Olito and Abbott (2025) (see their Appendix C and Fig. S5) report that invading X-linked inversions capturing the SDR-homologue and the female-beneficial allele at a sexually antagonistic locus never fix (including for equivalent parameter values to those considered here), but instead reach a stable polymorphism maintained by negative frequency-dependent selection. Based on the low equilibrium frequencies they observe (typically < 0.2; Olito and Abbott, 2025, Fig. S5 C-D), the authors argue that such suppressors are likely to be lost over time and thus unlikely to contribute meaningfully to the evolution of recombination suppression. The discrepancy between our results and those of Olito and Abbott (2025) arises from two mistakes in the population genetic recursions used to find equilibrium frequencies for an inverted genotype in Olito and Abbott (2025) (Colin Olito personal communication). This paper can thus be considered an update to Olito and Abbott (2025) with respect to the substitution of X-linked inversions.

Finally, balanced polymorphisms for X-linked inversions, such that invasion does not imply fixation, have been found for some specific parameter combinations (Blackmon and Brandvain, 2017). This occurs when A and a are strictly male/female-beneficial alleles (no overshooting) and the male-beneficial allele (A) is recessive in both sexes (i.e., female-beneficial allele a dominant) (e.g., Blackmon and Brandvain, 2017, Figure 1). However such alleles are expected to contribute weakly to sexually antagonistic variation and require non additive gene action phenotype (Connallon and Chenoweth, 2019; Fry, 2010; Jordan and Charlesworth, 2012), so we do not consider them here.

### D.3 Invasion of X-, Z- and W-suppressors under polygyny

Calculating the invasion probability of an X-linked suppressor from a multitype branching process in the presence of lottery polygyny is complex as it involves considering male and female fitness distributions (the latter of which depend on the distribution of mating success across males). We therefore estimated its fixation probability directly using individual-based simulations. This involved an identical simulation procedure as described in Appendix C.3.3, except that here a recombination suppressor was introduced as a single copy into a randomly sampled X-chromosome (rather than Y-chromosome). We then calculated the fixation probability for a given parameter set as the proportion of replicates in which the suppressor fixed. We used this proportion in place of Ψ^X^ when producing Fig. 4B.

To model the invasion of Z- and W-linked suppressors in an ZW system under polygyny, we followed a near-identical procedure as for XY simulations. The only difference here was that the SDR was now female-determining and found on the maternally inherited chromosome (W-chromosome). Suppressors then introduced on randomly sampled Z or W chromosome depending on treatment.

## E Simulations of deleterious variation

In this Appendix we outline our simulation procedure for the evolution of recombination suppression in the presence of deleterious variation distributed across many loci in the PAR. Our approach follows that of Roze and Lenormand (2025) but with the addition of a sexually antagonistic locus. Specifically we consider a chromosomal region containing *L* ≫ 1 deleterious loci, an SDR (modelled as a single locus), a sexually antagonistic locus, and a recombination modifier locus, giving a total of *L +* 3 loci. These loci are distributed as follows: deleterious loci are found at sites 1,…, *L*/2 − 1, *L*/2 *+* 1,…, *S* − 1, *S +* 1,…, *L +* 2, where *S* is the position of the sexually antagonistic locus, while the SDR and modifier loci are at sites *L*/2 and *L +* 3, respectively. Recombination amongst adjacent sites is assumed to be uniform across the region, occurring at a rate *r* ^°^. We are therefore considering a setup where an SDR is flanked by equal numbers (*L*/2) of deleterious loci to its left and right, with a sexually antagonistic locus nestled amongst the deleterious sites and whose position can be adjusted to modulate the effective recombination rate between this locus and the SDR (corresponding to the parameter *r* in our other models). The modifier locus occupies the terminal position and so (in the absence of a suppression mutation) does not influence recombination amongst the other loci.

During meiosis, the mean number of crossovers in our focal region is given by the map length *R* (unless otherwise stated, we assume *R =* 1), such that the genetic distance between adjacent sites is *R*^°^ *= R*/(*L +* 2). Because *R* is not large and we consider many deleterious loci, *R*^°^ ≪ 1 and so closely approximates the recombination rate amongst sites, denoted *r* ^°^. We therefore assume *r* ^°^ *= R*^°^ in our simulations. According to Haldane’s mapping function (Haldane, 1919), for the sexually antagonistic locus and the SDR to recombine with probability *r*, they must be separated by a genetic distance Log(1/(1 − 2*r*)/2. This entails that a number *d* (*r*) *=* Log(1/(1 − 2*r*))/(2*R*^°^) of sites sit between these two loci (we round *d* (*r*) to the nearest integer), which, given the SDR is always found at position *L*/2 means that to achieve a recombination rate *r* we must place the sexually antagonistic locus at position *S = L*/2 *+d* (*r*).

In the presence of deleterious variation, individuals’ fitness now depends on their expression of trait *z* and on the alleles they carry at deleterious loci. We assume all deleterious loci are di-allelic, with a wild-type allele that does not impact fitness and a deleterious allele that reduces fecundity. Mutations is unidirectional from the wild-type to the deleterious allele at a rate *µ*_*v*_ *=* 10^−4^. The costs of deleterious alleles combine multiplicatively across loci such that a male’s and female’s fecundity is given by

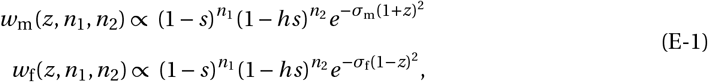

where *n*_1_ and *n*_2_ are the number of loci for which an individual is heterozygous and homozygous respectively, *s* is the homozygous selection coefficient of a deleterious allele, and *h* is the dominance coefficient.

We ran simulations for different parameter combinations of selection *s* (0.005, 0.025, 0.125), dominance *h* (0.1, 0.5) coefficients and for the numbers of deleterious loci *L* (100, 500, 1000), and did so for an XY system, where the SDR is male-determining and paternally inherited, and a ZY system, where the SDR is female-determining and maternally inherited. For each parameter combination we simulated 10 replicate burn-in populations (in the absence of recombination modifiers and without sexually antagonistic selection: *σ =* 0) for 5000 generations, at which point allele frequencies at the deleterious loci had reached mutation-selection-drift equilibrium. We introduced allele A at the sexually antagonistic locus at its equilibrium frequencies in all gene pools (i.e. for XY system 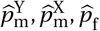, see Appendix A.2.1 for calculations of these frequencies). Then, we let a single recombination suppressor arise, one that captures an entire haploid chromosome (all *L +* 3 loci) at the same time. This suppressor appeared on a randomly sampled X, Y, Z, or W chromosome, depending on which type of modifier was being studied, and we tracked its frequency until it fixed or was lost. We repeated this procedure 10^6^ times for each burn-in population, giving 10 estimates of the fixation probability for each parameter combination. Where relevant, all other aspects of the simulation proceed as in Appendix C.3.2.

---

**Supplementary Figure 1.**
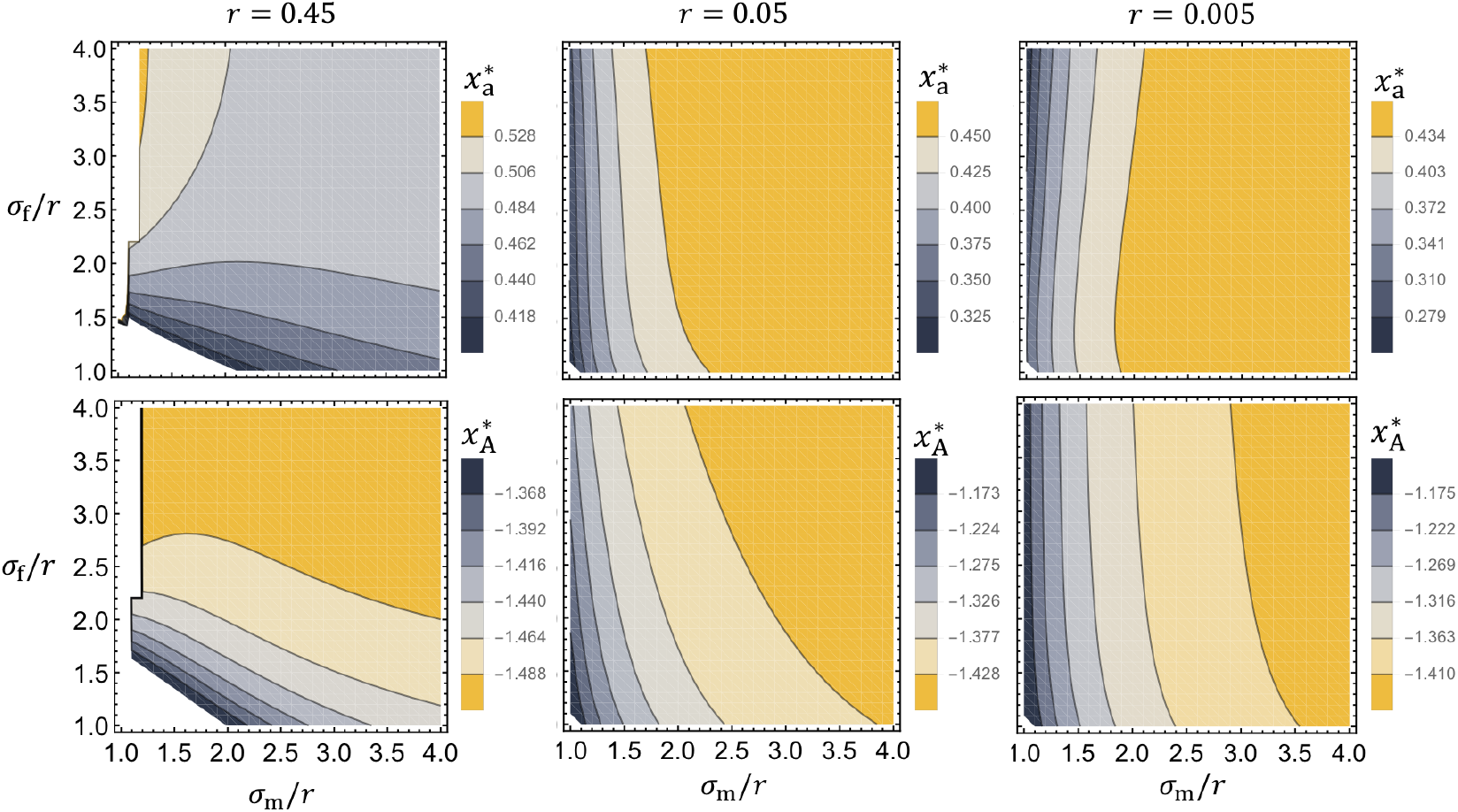
Allelic values at the evolutionary stable values 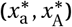. Top row panels show the allelic value for 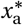 at the stable values 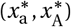 (Appendix A for calculation) as a function of the strength of male- and female-selection at the trait locus relative to the recombination rate (*σ*_m_/*r* and *σ*_f_/*r*). Bottom rows show equivalent results for the allelic value 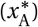. Row panels show results for different strengths of recombination between the trait locus and SDR (*r*). Non-coloured space corresponds to where evolutionary branching at the trait locus does not occur (condition (A-2) not fulfilled). As seen under strictly symmetric sex-specific selection (*σ = σ*_m_ *= σ*_f_, Fig. 1C), 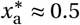 and *x*_A_ ≈ −1.5 unless sexually antagonistic selection is weak relative to the strength of recombination (*σ*_m_ and *σ*_f_ small while *r* big).

**Supplementary Figure 2.**
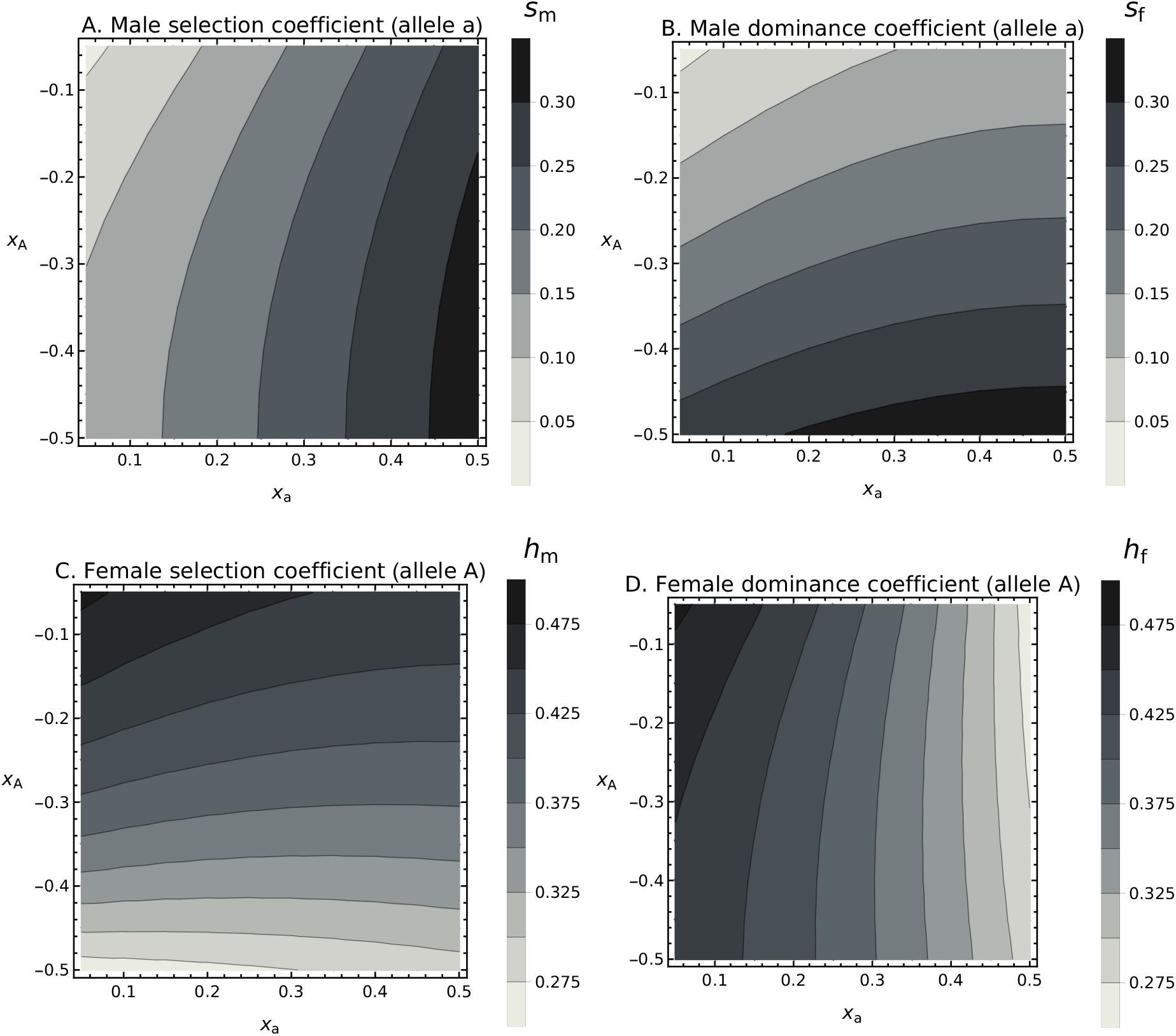
Selection and dominance coefficients in convention of Kidwell et al. **(1977)**. Panels **A**. and **B**. show male (*s*_m_) and female (*s*_f_) selection coefficients, defined as the relative fitness cost when homozygote for the non-preferred allele: allele a imposes a cost *s*_m_ *=* [*w*_m_(2*x*_A_) −*w*_m_(2*x*_a_)]/*w*_m_(2*x*_A_) in males, and alleles A imposes a cost *s*_f_ *=* [*w*_f_(2*x*_a_) −*w*_f_(2*x*_A_)]/*w*_f_(2*x*_a_) in females. Panels **C**. and **D**. show the male (*h*_m_) and female (*h*_f_) dominance coefficients: *h*_m_ *=* [*w*_m_(2*x*_A_) − *w*_m_(*x*_a_ *+ x*_A_)]/[*w*_m_(2*x*_A_) − *w*_m_(2*x*_a_)] and *h*_f_ *=* [*w*_f_(2*x*_a_) − *w*_f_(*x*_a_ *+ x*_A_)]/[*w*_f_(2*x*_a_) − *w*_f_(2*x*_A_)]. Values are plotted here for *σ = σ*_m_ *= σ*_f_ *=* 0.1 as in main text Fig. 2.

**Supplementary Figure 3.**
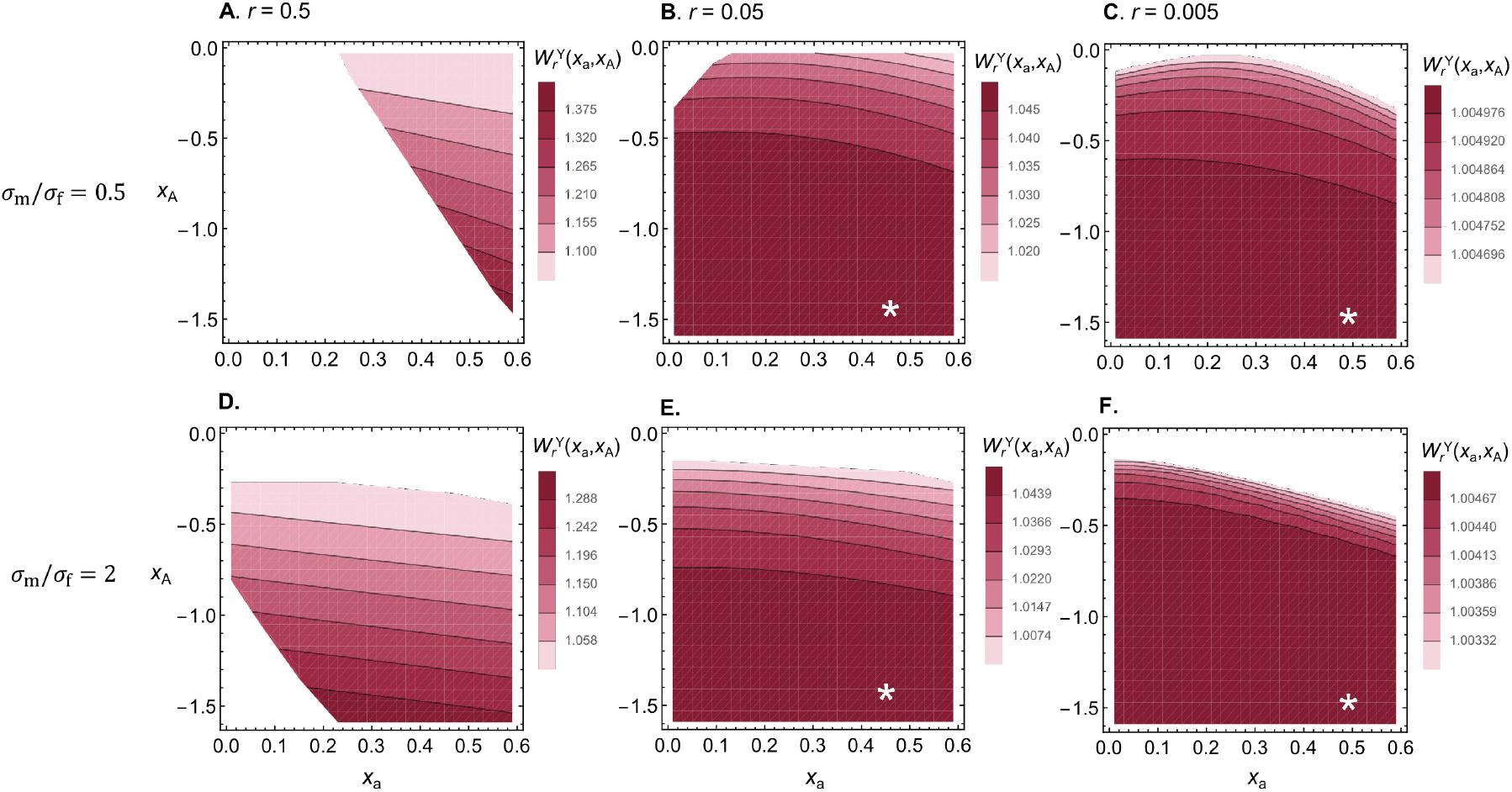
Invasion fitness of a Y-linked suppressor capturing a trait locus with alleles encoding *x*_a_ and *x*_A_ under sex-biased selection. Plots show the invasion fitness, 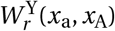 (main text eq. C-4, Appendix C.1 for calculation details), of a Y-linked recombination suppressor capturing allele A at the trait locus as a function of the allelic values *x*_a_ and *x*_A_. Panels **A-C** (top row) show invasion fitness in the presence of female-biased selection at the trait locus (*σ*_m_ *=* 0.075, *σ*_f_ *=* 0.15, so that *σ*_m_/*σ*_f_ *=* 0.5), and panels **D-F** show results for male-biased selection (*σ*_m_ *=* 0.15, *σ*_f_ *=* 0.075, so that *σ*_m_/*σ*_f_ *=* 2). Different row panels show results for different levels of resident recombination between the trait locus and the SDR, *r*. White star shows the location of the evolutionary stable allelic values 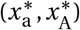. Compared to the case of equally strong selection in both sexes (see Fig. 2), the conditions for a Y-linked suppressor to invade are more restrictive under sex-biased selection, which is because such selection is less amenable to the maintenance of a balanced polymorphism at the trait locus (i.e., maintains polymorphism for a narrower range of allelic value combinations; *x*_a_ and *x*_A_).

**Supplementary Figure 4.**
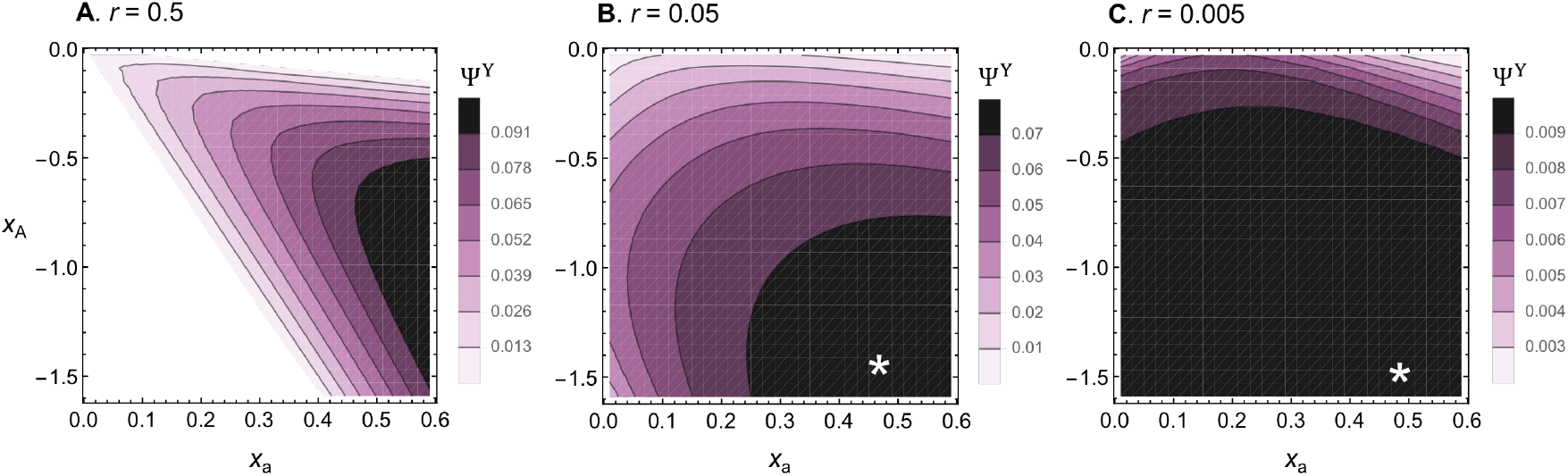
Invasion probability of a Y-linked suppressor capturing a trait locus with alleles encoding *x*_a_ and *x*_A_. Plots show the invasion probability under the random-union-of-gametes model, Ψ^Y^ (Appendix C.3.1 for calculation details), of a Y-linked recombination suppressor capturing allele A at the trait locus as a function of the allelic values *x*_a_ and *x*_A_. Different panels show results for different levels of resident recombination between the trait locus and the SDR, *r*. White star shows the location of the evolutionary stable allelic values 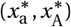. Other parameters used *σ = σ*_m_ *= σ*_f_ *=* 0.1. See SI Fig. 2 for the selection and dominance coefficients of alleles A and a in the notation of Kidwell et al. (1977).

**Supplementary Figure 5.**
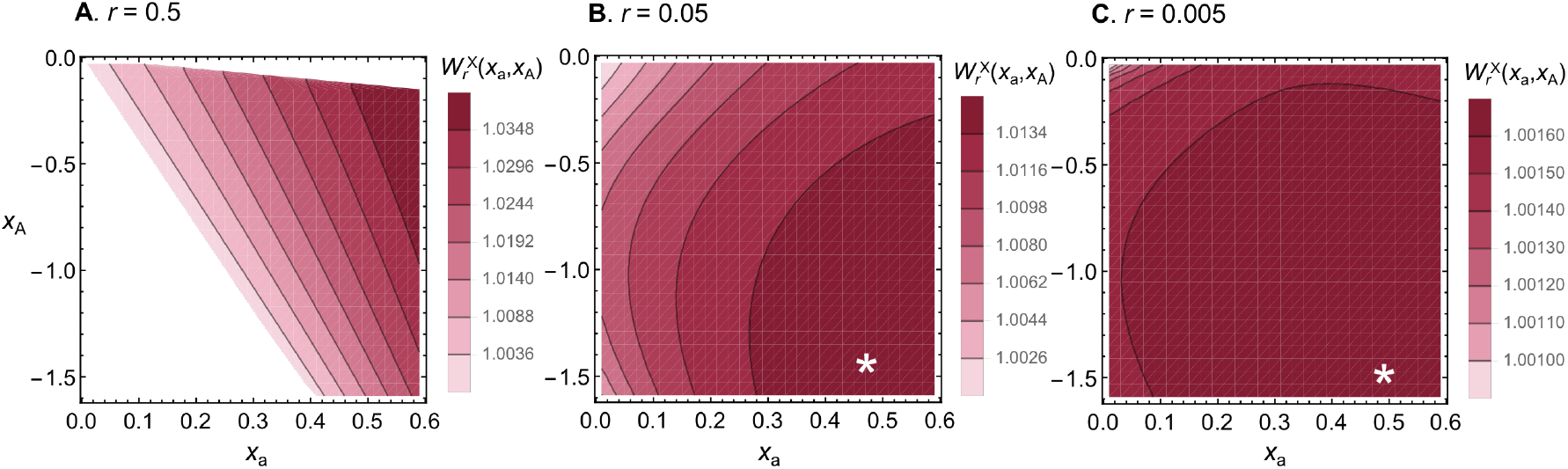
Invasion fitness of an X-linked suppressor capturing a trait locus with alleles encoding *x*_a_ and *x*_A_. Plots show the invasion fitness, 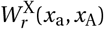 (main text eq. C-4, Appendix D.1 for calculation details), of an X-linked recombination suppressor capturing allele a at the trait locus as a function of the allelic values *x*_a_ and *x*_A_. Different panels show results for different levels of resident recombination between the trait locus and the SDR, *r*. White stars shows the location of the evolutionary stable allelic values 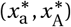. Other parameters used *σ = σ*_m_ *= σ*_f_ *=* 0.1. See SI Fig. 2 for the selection and dominance coefficients of alleles A and a in the notation of Kidwell et al. (1977).

**Supplementary Figure 6.**
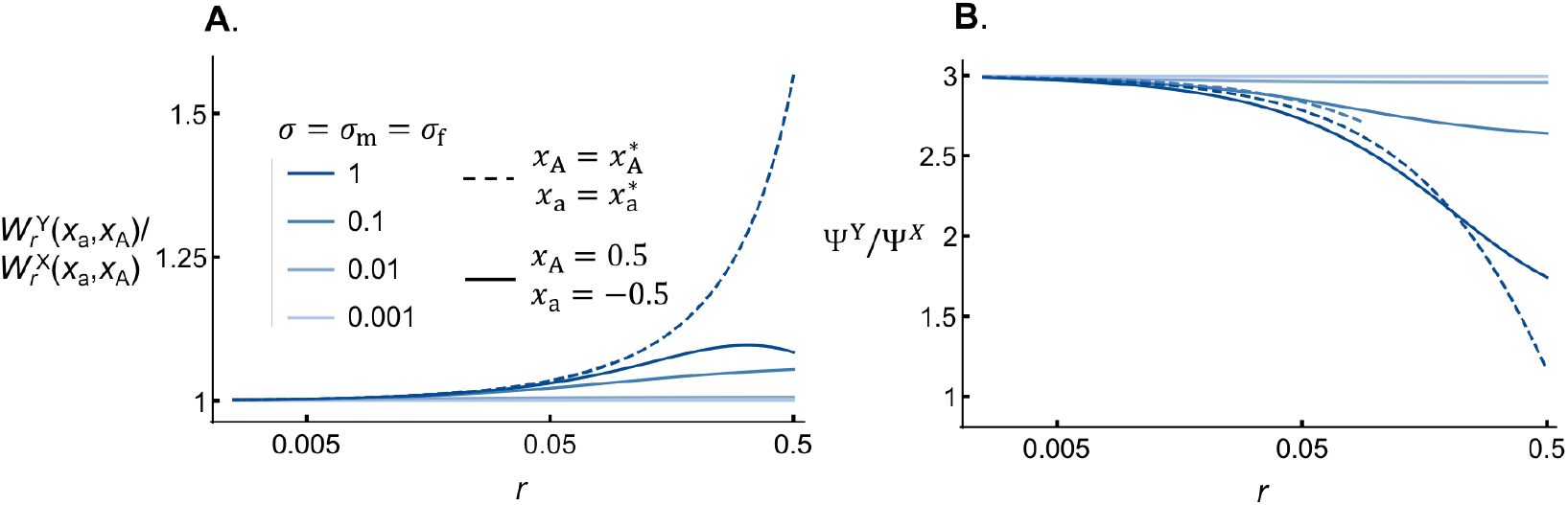
Invasion fitness and invasion probability for Y- vs X-linked suppressors. Panel **A**. shows the ratio of the invasion fitnesses 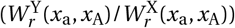 of a Y-linked suppressor capturing allele A at the trait locus and an X-linked suppressor capturing allele a for different recombination rates *r* and strength of sexually antagonistic selection *σ = σ*_m_ *= σ*_f_. Dashed lines correspond to results for evolutionary stable allelic values 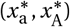 and solid lines for *x*_a_ *=* 0.5, *x*_A_ *=* −0.5 (Appendices C.1 and D.1 for calculation of 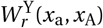 and 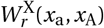). Panel **B**. shows the ratio of the Y- vs X-linked invasion probabilities (Ψ^Y^/Ψ^X^) under the random-union-of-gametes model for the same parameter combinations and using the same legend as in panel A (Appendices C.3.1 and D.2.1 for calculation of Ψ^Y^ and Ψ^X^).

**Supplementary Figure 7.**
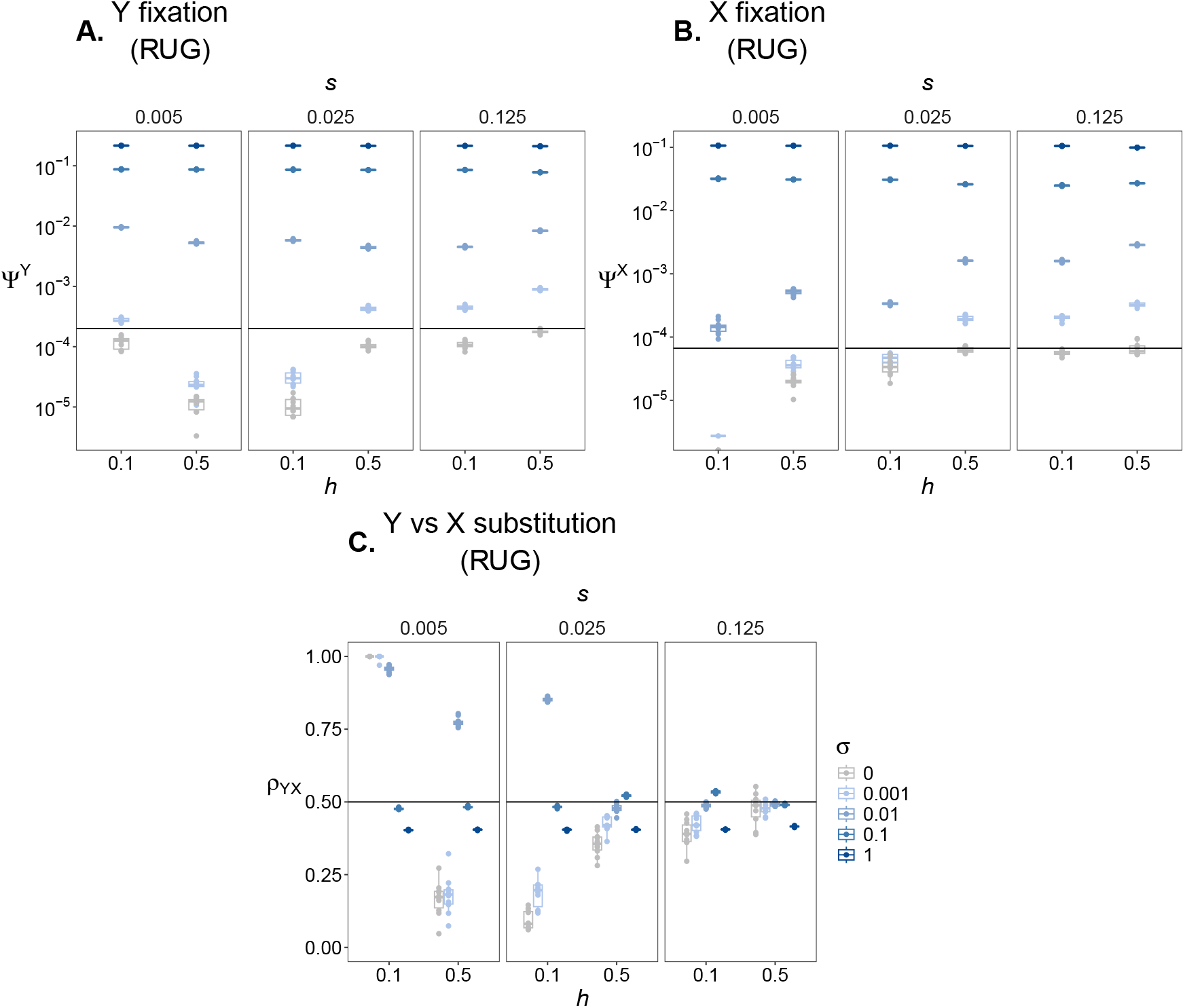
Selection for recombination suppression when deleterious and sexually antagonistic alleles co-segregate. Panel **A**. shows the fixation probabilities estimated from individual-based-simulations of a Y-linked suppressor capturing both a sexually antagonistic locus and *L =* 100 deleterious loci at mutation-selection balance (Appendix E for simulation details). Results are shown for selection coefficient *s =* 0.005, *s =* 0.025, and *s =* 0.125 (see columns) and dominance coefficients of either *h =* 0.1 and *h =* 0.5 (see x-axis) at all deleterious loci. Each dot is an estimate for a single replicate population in a simulation of the random union-of-gametes model. Boxplot and dot colours correspond to different strengths of sexually antagonistic selection (*σ*). Horizontal line is the neutral fixation probability for a Y-linked mutation (1/(*N* /2)), where *N =* 10^4^. Recombination is scaled such that *r =* 0.3 between the sexually antagonistic locus and SDR (Appendix E). Panel **B**. shows equivalent results as panel A, but for X-linked suppressors (here the neutral probability is 1/(3*N* /2)). Panel **C**. shows the probability that a substitution by a recombination suppressor is Y- vs. X-linked, *ρ*_YX_, from the same simulation setup as in panel A. Horizontal line marks *ρ*_YX_ *=* 0.5. Here, XY and ZW systems are interchangeable: Ψ^Y^ *=* Ψ^W^, Ψ^X^ *=* Ψ^Z^, and *ρ*_YX_ *= ρ*_WZ_.

